# Fatty acids in the tumor microenvironment reprogram neutrophils to induce immunosuppression via adenosine

**DOI:** 10.64898/2026.04.02.716169

**Authors:** Rashi Singhal, Nina W. Zhang, Zheng Hong Lee, Hannah N. Bell, Prarthana J. Dalal, Sumeet Solanki, Wesley Huang, Ryan Rebernick, Peter Sajjakulnukit, Himani Jasewicz, Roshan Kumar, Nikhil K. Kotla, Amanda Huber, Anitha Vijay Mallappa, Subhash B. Arya, Shogo Takahashi, Carlos Ichiro Kasano-Camones, Eileen Carpenter, Marina Pasca di Magliano, James J. Moon, Carole A. Parent, Frank J Gonzalez, Andrew D. Patterson, Michael D. Green, Weiping Zou, Elena M. Stoffel, Costas A. Lyssiotis, Yatrik M. Shah

## Abstract

As solid tumors progress, the tumor microenvironment (TME) becomes increasingly immunosuppressive, impairing cytotoxic T-cell activity and limiting the efficacy of the immune checkpoint blockade. However, the mechanistic drivers of this immunosuppression remain poorly understood. Here, we identify a tumor-derived lipid–neutrophil–adenosine axis as a critical regulator of immune suppression in advanced colorectal cancer (CRC). We show that fatty acids enriched in tumor interstitial fluid reprogram neutrophils to generate adenosine via PPARα activation, leading to T-cell suppression. Using AB928, a dual A2aR/A2bR adenosine receptor antagonist currently in clinical trials, we restored T-cell proliferation, effector function, and tumor-killing capacity in vitro and in vivo. Importantly, AB928 synergized with anti-PD-1 therapy to enhance survival in an autochthonous model of metastatic CRC. Our findings define a metabolic immune evasion mechanism in the TME and provide a rationale for targeting neutrophil-derived adenosine signaling to improve immunotherapy responses in CRC and other solid tumors.

## Introduction

The tumor microenvironment (TME) is prone to frequent metabolic changes that reprogram immune cell function and affect antitumor immunity (1). Neutrophils are a major immune cell type in the TME that displays considerable heterogeneity (2). They are frequently associated with poor outcomes in cancer patients, and increased tumor infiltration correlates with worse prognosis (3–6). However, neutrophils can also exert antitumor effects, either through direct cytotoxicity or via interactions with other immune cells (7–10). Their role in cancer remains controversial (11); while some studies report reduced tumor burden following anti-Ly6G–mediated neutrophil depletion, others demonstrate tumor-restraining effects of neutrophils (12–15). In early-stage tumors, patients responding to immunotherapy exhibit a distinct subset of neutrophils capable of controlling tumor growth (16,17). These findings highlight the plasticity of neutrophils, which can adopt either pro- or antitumor phenotypes depending on microenvironmental cues (18).

Understanding how metabolism influences neutrophil function in the TME is critical. Alongside macrophages, neutrophils are considered immunosuppressive myeloid cells that contribute to poor responses to immune checkpoint blockade (ICB) (19–26). While cytokines and chemokines are known to shape neutrophil behavior, the impact of nutrient availability remains poorly defined. Cancer cell proliferation and effective immune responses both rely on access to nutrients within the TME (27,28). Lipid accumulation is a hallmark of many solid tumors and promotes tumor progression (29–31). Additionally, lipid metabolism also plays a central role in suppressing antitumor immunity (32,33). The tumor interstitial fluid (TIF), which represents the extracellular medium of the TME, in a B16 melanoma tumor model in mice contains many fatty acids (FAs), the abundance of which increases as tumors progress (34). Numerous studies have focused on the effects of fatty acids on T cells and have mostly concentrated on showing the toxic effects of fatty acids at high concentrations via the induction of several pathways (29,35,36). Studies have shown that several lipid species can exhibit signaling effects on the differentiation and polarization of tumor-associated macrophages and play a significant role in inducing immunosuppressive effects that support tumor growth and progression (37–41).

Despite these advances, how lipids shape the immune landscape, particularly neutrophil function remains unclear. Here, we show that neutrophils in advanced tumors adopt a highly suppressive phenotype compared to those in early tumors. This transition is driven, in part, by nutrient signals within the TIF. FAs promote the emergence of immunosuppressive neutrophils that produce adenosine, a potent inhibitor of T cell proliferation and activation. Multiple FA species were sufficient to induce neutrophil-mediated adenosine release. Blocking the adenosine receptor on T cells with a selective antagonist reversed the suppressive effects of FA-educated neutrophils and improved survival in an immunosuppressive colorectal cancer (CRC) mouse model when combined with immunotherapy. These findings identify a fatty acid–neutrophil–adenosine axis as a selective and targetable pathway to restore T-cell cytotoxicity and enhance antitumor immunity in the TME.

## Methods

### Mice and treatments

Wild-type (WT) C57BL/6 or BALB/C mice were used for syngeneic experiments. *TripleMut* and PPARα KO mice were described previously(42,43). OT-1 mice were from Jackson laboratories (strain #:003831) and were used for *in vitro* or *in vivo* T-cell killing assays. Mice were kept in individually ventilated cages under specific pathogen-free conditions at a room temperature of 22°C ± 2°C, 45–65% relative humidity, and a light/dark cycle (12 h of light/12 h of dark). Food and drinking water were provided ad libitum with a standard chow diet. Six- to eight-week-old male or female mice were used for all experiments. All animal studies were carried out in accordance with the Association for Assessment and Accreditation of Laboratory Animal Care International guidelines and approved by the University Committee on the Use and Care of Animals at the University of Michigan and the National Cancer Institute Animal Care and Use Committee under Protocol LM-016. For the syngeneic model, 2 million MC38 or Yumm5.2 and 1 million CT26 cells were subcutaneously injected into both flanks of C57BL/6 or BALB/c mice, respectively. Tumor size was measured with digital calipers. At 30 days (MC38 model) or 20 days (CT26 and Yumm5.2 model) after the tumor growth reached the advanced stage, the mice were euthanized, and the neutrophils were isolated. *TripleMut* mice were induced with tamoxifen (Sigma-Aldrich) dissolved in corn oil at either 50 mg/kg or 100 mg/kg administered by intraperitoneal injection for 3 consecutive days. For survival studies, mice received anti-PD-1 antibody (clone RMP1-14, Bio X Cell) at 5 mg/kg intraperitoneally every 3 days starting 3 days post-tamoxifen induction. Anti-Ly6G antibody (clone 1A8, Bio X Cell) was administered at 100 μg per mouse intraperitoneally every 3 days. Anti-CD8α antibody (clone 2.43, Bio X Cell) was given at 25 mg/kg intraperitoneally every 5 days. Control groups received isotype-matched control antibodies (IgG2a for anti-PD-1, IgG2b for anti-Ly6G and anti-CD8α) at equivalent doses and schedules.

### Tumor enteroid culture and in vivo growth

The patient-derived 14481 enteroid line, harboring mutations in APC, KRAS, and TP53, was cultured as previously described(44,45). Enteroids were dissociated into single cells, resuspended in 200 µL of Matrigel (Corning), and injected subcutaneously into the flanks of nude mice, which lack T cells but retain functional innate immune populations.

### Cell lines

The MC38, MC38-Ova, and CT26 mouse colon carcinoma cell lines were maintained in RPMI-1640 medium and the Yumm5.2 mouse melanoma cell line were maintained in DMEM F12 medium supplemented with 10% fetal bovine serum and 1% antibiotic/antimycotic. All cell lines were regularly tested for possible mycoplasma contamination. All cell lines used in this study were mycoplasma negative.

### Primary cell cultures

Bone marrow-derived neutrophils were purified from bone marrow cells isolated from the femurs and tibias of 6- to 8-week-old WT mice or tumor-bearing mice (C57Bl6/J) using a two-layer density gradient method with 62.5% Ficoll (GE Healthcare) as a separation reagent in 1X HBSS-Prep (HBSS + 0.5% FBS + 20 mM Na-HEPES (pH 7.4) + Pen/Strep). Bone marrow cells were layered over 62.5% Percoll and centrifuged for 30 min at 1000 × g with no brake and no acceleration. At the end of the gradient centrifugation, a sharp interface on top of the 62.5% percoll (these are immature cells and non-granulocytic lineages) was separated from a cloudier pellet (the neutrophils). The neutrophil pellet was washed in HBSS prep, and the cells were cultured for 16-24 hours in RPMI with GlutaMax + 10% heat-inactivated FBS + 20 mM HEPES + and 1% penicillin/streptomycin supplemented with 10 ng/mL GM-CSF (BioLegend). Tumor neutrophils were isolated from tumors harvested from syngeneic model mice. The tumors were washed, dissected into small pieces, and digested with collagenase V for 45 minutes at 37°C. The digested pieces were then collected, and a single-cell suspension was made. Neutrophils were extracted from single-cell suspensions of tumors using the MojoSort™ Mouse Neutrophil Isolation Kit (BioLegend) according to the manufacturer’s instructions. For colon neutrophils, immune cells were first enriched and separated from epithelial cells, for which the colon was washed twice with PBS and cut into small pieces, followed by digestion with 25 mM EDTA twice in a 37°C incubator shaker to remove epithelial cells. This step was followed by digestion with 0.5 mg/mL collagenase IV, after which the cells were purified by using a 40%–70% Percoll gradient. The immune cell extract was then subjected to neutrophil isolation using the MojoSort™ Mouse Neutrophil Isolation Kit (BioLegend) according to the manufacturer’s instructions. For the *in vitro* proliferation or activation assay, CD3+ T or CD8+ T cells were isolated from the splenocytes of 6- to 8-week-old C57Bl6/J mice using a mouse MojoSort mouse CD3+ T or CD8+ T-cell isolation kit (BioLegend) according to the manufacturer’s instructions. After CD3+ or CD8+ T-cell isolation, 1 × 10^5^ splenocytes per well were seeded on αCD3 (1.0 μg/ml, clone: 145-2C11, Thermo Fisher) and αCD28 (2 μg/ml clone: 37.51, Thermo Fisher) -coated 96-well round-bottom plates in RPMI-1640 (Gibco) supplemented with 10% FBS, 1% penicillin/streptomycin, 1% L-glutamine, 10 mM HEPES, 1% nonessential amino acids, 1% sodium pyruvate, 0.1% β-mercaptoethanol and 5 ng/mL IL2. For co-culture experiments, neutrophils or macrophages treated with the indicated reagents in the figure legends or left untreated were cocultured with CD3+ T cells at a ratio of 1:2.

### Adenosine receptor and β-oxidation inhibitor treatments

AB928 (Arcus Biosciences) and AB680 (Arcus Biosciences) were dissolved in DMSO to prepare 10 mM stock solutions and stored at -20°C. For in vitro experiments, T cells were pretreated with 1 μM AB928 or cancer cells were pretreated with 100 nM AB680 for 30 minutes at 37°C prior to coculture. For in vivo experiments, AB928 was dissolved in 0.5% methylcellulose and administered by oral gavage at 100 mg/kg twice daily. Etomoxir (Sigma-Aldrich) was dissolved in DMSO and used at concentrations of 10 μM or 25 μM as indicated in figure legends with 16-hour pretreatment prior to functional assays.

### Human tissue collection

Fresh human colorectal adenoma and adenocarcinoma tissue samples were obtained from patients undergoing surgical resection at the University of Michigan Medical Center under protocols approved by the Institutional Review Board (IRB protocol #HUM00064405). Written informed consent was obtained from all patients prior to tissue collection.

### Neutrophil isolation

Tumor tissues were minced into 2-3 mm pieces and digested with 1 mg/mL collagenase IV (Worthington Biochemical) and 0.1 mg/mL DNase I (Sigma-Aldrich) in HBSS for 45 minutes at 37°C with gentle agitation. Single-cell suspensions were filtered through a 70 μm cell strainer and resuspended in PBS. Human neutrophils were isolated using a discontinuous density gradient with Histopaque-1077 and Histopaque-1119 (Sigma-Aldrich). Briefly, 3 mL of Histopaque-1119 was layered beneath 3 mL of Histopaque-1077 in a 15 mL conical tube. The cell suspension (up to 6 mL) was carefully layered on top of the Histopaque-1077 and centrifuged at 700 × g for 30 minutes at room temperature with no brake. Neutrophils were collected from the interface between Histopaque-1119 and the cell pellet, washed twice with HBSS, and resuspended in complete RPMI medium. Neutrophil purity was confirmed to be >90% by flow cytometry using CD66b and CD15 surface markers, and viability was assessed by trypan blue exclusion to be >95%.

### Tumor proliferation and apoptosis quantification

For proliferation analysis, mice were injected intraperitoneally with 100 mg/kg BrdU (Sigma-Aldrich) 2 hours prior to sacrifice. Tumor tissues were fixed in 4% paraformaldehyde, paraffin-embedded, and sectioned at 5 μm thickness. BrdU incorporation was detected using the BrdU In-Situ Detection Kit (BD Pharmingen) according to the manufacturer’s protocol. Apoptosis was assessed using the In Situ Cell Death Detection Kit (Roche) following standard TUNEL staining procedures. Proliferation was quantified as the percentage of BrdU-positive cells, and apoptosis was quantified as the percentage of TUNEL-positive cells in at least 5 random high-power fields (40× magnification) per tumor section using ImageJ software.

### NETosis assay

NETosis was quantified using a real-time fluorescence imaging approach coupled with automated cell segmentation and classification. Bone marrow-derived neutrophils were seeded at 2 × 10L cells per well in 8-well glass-bottom chamber slides (Cellvis, #C8-1.5H-N), precoated with 25 μg/ml fibrinogen and 10 μg/ml fibronectin in DPBS. Cells were cultured overnight at 37°C in 5% CO₂ with either oleic acid (100 μM), arachidonic acid (100 nM), or BSA (vehicle control) in RPMI-1640 supplemented with GlutaMAX, 10% heat-inactivated FBS, 20 mM HEPES, 1% penicillin/streptomycin, and 10 ng/ml GM-CSF.

The following day, the media was replaced with RPMI containing 2% BSA. Cells were stained with Hoechst 33342 (1 μg/ml), CellMask Orange (1 μM), and SYTOX Green (500 nM), and stimulated with 100 nM phorbol 12-myristate 13-acetate (PMA; Sigma-Aldrich) to induce NETosis. Images were acquired every minute for 2 hours using a Zeiss Colibri fluorescence microscope at 20× magnification. Image analysis was performed using CellProfiler, where nuclei and cell boundaries were segmented, and NETotic cells were identified as those with SYTOX Green-positive cytoplasm. NETosis was quantified as the percentage of NETotic cells over the total cell count at 90 minutes post-PMA treatment.

### T-cell proliferation and activation assay

T cells were stained prior to seeding with 2 μM CFSE (Biolegend, San Diego, CA, USA) dye for 15 min at 37°C, according to the manufacturer’s instructions. Excess dye was removed by washing three times with CFSE staining buffer (PBS+1% FBS). T cells were cocultured with neutrophils previously treated or not treated with different reagents. Three days later, the cells were stained with antibodies against CD4, CD8, CD44 or CD25 (Invitrogen) for specific subset gating and with 7AAD (BD Biosciences) for dead cell exclusion. Single-cell suspensions were taken for flow cytometry analysis. Since CFSE is diluted by approximately half with each cell division, live single cells (negative 7AAD staining) were gated, and the percent proliferation was calculated by gating the dim CFSE (left) population. The mean fluorescence intensity of CD25 and the percentage of CD44-positive cells was used to evaluate the level of CD4+ and CD8+ T-cell activation.

### T-cell exhaustion and intracellular cytokine assays

CD8+ T cells were treated with PMA, ionomycin, and brefeldin A (Sigma) for activation and GolgiStop (BD Pharmingen) to stop extracellular release at 6 hours prior to the assay. The cells were first stained for surface markers by using antibodies against CD8, PD-1 and Tim-3 (eBioscience) and the live-dead exclusion marker 7AAD (BD Biosciences), after which they were washed twice with FACS buffer. The cells were then incubated in fixation/perm buffer (eBioscience #005123) overnight at 4°C. The next day, the cells were first washed twice with permeabilization buffer (eBioscience #008333), and then CD8+ T-cell cytokine expression was determined by intracellular staining with Abs against mouse IFN-γ, TNF-α and granzyme-b (Invitrogen) prepared in permeabilization buffer for 45 minutes on ice. The samples were washed thrice with permeabilization buffer, and all flow samples were resuspended in FACS buffer and acquired through a Cytek Aurora Flow cytometer with a minimum of 50,000 events captured per sample.

### *In vitro* and *in vivo* cytotoxic CD8+ T-cell killing assay

MC38 or B16 cells expressing ovalbumin were suspended in complete RPMI 1640 medium (as described for the T-cell culture above). Cells were seeded in a 24-well plate at 50,000 cells per well and incubated for 24 h to enable their attachment to the substrate. CD8+ T cells were isolated from OT-1 transgenic mouse splenocytes using a CD8+ MojoSort negative selection kit (BioLegend). CD8+ T cells were activated with 5 μg/ml ovalbumin peptide (OVA257-264, InvivoGen, San Diego, CA, USA) and cultured for 3 or 4 days in complete RPMI-T-cell medium with or without neutrophil-pretreated conditioned media. They were then added on top of the MC38 Ova cells. Then, 50 ng/mL SYTOX Green dye (Invitrogen) was added, and the entire plate was imaged via live cell imaging using a Cytation 5 Imaging Multi-Mode reader to quantify the green-fluorescent cells. Alternatively, cells were stained with 7AAD for cell death staining and analyzed on a flow cytometer (Cytek Aurora Spectral Analyzer). The percentage of cell death was calculated using cells positively stained for 7AAD. For the *in vivo* assay, CD8+ T cells isolated from 10 OT-1 mice were cultured for 4 days in conditioned medium from neutrophils induced with OA in the presence of AB928, after which they were counted, and 1 million cells were adoptively transferred into individual MC38-OVA-bearing mice. Tumor progression was monitored throughout the study, and the terminal weights of the tumors were recorded for each group. Proliferation and apoptosis were evaluated in tumor cells

### Tumor Interstitial Fluid (TIF) and Serum Isolation

Mouse tumor interstitial fluid (TIF) was isolated from MC38, CT26 or Yumm5.2 tumors using a protocol modified from a previously described centrifugation method (34,46–49). Tumor-bearing mice were euthanized by cervical dislocation, and tumors were rapidly dissected from the animals within 1 min. Blood was collected from the same animal via cardiac puncture and centrifuged at 800 × g for 10 min at 4°C to separate the plasma. The plasma was frozen in liquid nitrogen and stored at -80°C until further analysis. Tumors weighing greater than 500 mg were quickly rinsed in 1X PBS, and extra PBS was added on filter paper. The tumors were then placed onto a 15 μm cell strainer, PluriStrainer (PluriSelect, CA, USA), fixed on top of 50 mL conical tubes and centrifuged at 150 × g at 4°C for 10 min. TIFs were then collected, transferred to a 1.5 mL tube, frozen and stored at -80°C until further analysis. TIF (30–100 μL) was isolated from 1 large tumor weighing greater than 500 mg.

Human TIF was isolated from fresh adenoma and adenocarcinoma tissues within 15 minutes of surgical resection. During transport, tissues were kept on ice to preserve integrity. Tumors weighing between 40–75 mg were placed onto 15 μm PluriStrainers (PluriSelect, CA, USA) fitted on top of 50 mL conical tubes and centrifuged at 1000 × g for 10 minutes at 4°C. The resulting TIF (3–5 μL per sample) was collected and stored at –80°C until further analysis.

### Metabolomics

For neutrophil samples, 3 million cells were seeded in complete RPMI in a 6-well plate and treated with fatty acids or left untreated for early- or late-stage tumor neutrophils for 24 hours. The samples were centrifuged, and the supernatants were collected in separate tubes and immediately placed on dry ice. Two hundred microliters of neutrophil supernatant were mixed with 800 μL of ice-cold HPLC-grade 100% methanol. The neutrophil pellet was washed once with ice-cold PBS and then incubated in dry-ice cold 80% methanol on dry ice for 10 min before homogenization. The concentrations of the samples were normalized by conducting a BCA protein assay on a technical replicate, and the volume was normalized across samples to achieve even concentrations. Metabolite extracts from the supernatant or the pellet fraction were then lyophilized using a SpeedVac concentrator at 4°C and resuspended in a 50:50 methanol/water mixture for LCLMS analysis. The data were collected in both negative and positive ion modes. For negative mode, previously published parameters were used (50,51). For positive mode, an Agilent Technologies Triple Quad 6470 LC/MS system consisting of a 1290 Infinity II LC Flexible Pump (Quaternary Pump), 1290 Infinity II Multi sampler, 1290 Infinity II Multicolumn Thermostat with 6 port valve and 6470 triple quad mass spectrometer was used.

Agilent Mass Hunter Workstation Software LC/MS Data Acquisition for 6400 Series Triple Quadrupole MS with Version B.08.02 was used for calibration, compound optimization and sample data acquisition. A Waters Acquity UPLC HSS T3 1.8 mmVanGuard Pre-Column 2.1 × 5 mm column and a Waters UPLC BEH TSS C18 column (2.1 × 100 mm, 1.7 mm) were used with mobile phase A) consisting of 0.1% formic acid in water and mobile phase (B) consisting of 0.1% formic acid in acetonitrile. The solvent gradient was as follows: mobile phase (B) was held at 0.1% for 3 min, increased to 6% at 12 min, 15% at 15 min, 99% at 17 min and held for 2 min before reverting to the initial conditions and held for 5 min. The column was held at 40°C, and 3 ml of sample was injected into the LCMS at a flow rate of 0.2 ml/min. Calibration of the 6470 QqQ MS instrument was achieved through an Agilent ESI-Low Concentration Tuning Mix. QQ data for both ion modes were preprocessed with Agilent MassHunter Workstation Quantitative Analysis Software (B0700). Metabolite chromatogram peaks were manually inspected. The abundance of each metabolite was median normalized across the sample population. Statistical significance was determined by one-way ANOVA with a significance threshold of 0.05. For TIF and serum samples, 100 μL of TIF from 3 different tumors from separate mice and 100 μL of serum were mixed with 400 μL of 100% MeOH. The samples were then dried and resuspended in 1:1 MeOH/H2O and further processed as described for neutrophils.

### Lipidomics

Samples were analyzed by a Vanquish UHPLC system coupled with an Orbitrap Fusion Lumos Tribrid™ mass spectrometer using an H-ESI™ ion source (all Thermo Fisher Scientific) with a Waters (Milford, MA) CSH C18 column (1.0L× 150 mm, 1.7 µm particle size) in positive and negative ion detection modes. Solvent A was ACN: H2O (60:40; v/v) containing 10 mM ammonium formate and 0.1% formic acid, and solvent B was IPA: ACN (95:5; v/v) containing 10 mM ammonium formate and 0.1% formic acid. The flow rate of the mobile phase was 0.11 mL/min, and the column temperature was 65°C. The gradient of solvent B was 0 min 15% (B), 1–3 min 30% (B), 3–3.5 min 48% (B), 3.5–12 min 82% (B), 12–12.2 min 99% (B), 12.2–17.00 min 99% (B), 17.00–17.50 min 15% (B), and 17.00–20.00 min 15% (B). The ion source spray voltages are 4,000 V and 3,000 V in positive and negative mode, respectively. Mass spectrometry was conducted with a scan range from 200 to 1500 m/z for full scan and utilized in AcquireX mode with a stepped collision energy of 35% and 15% spread for the fragment ion MS/MS scan.

### Single-cell RNA sequencing

#### Library preparation and processing

Approximately 5-6 million bone marrow-derived neutrophils were isolated from 5 late-stage MC38 tumor-bearing mice. Cells from each mouse were pooled and processed for single-cell RNA sequencing using an Chromium single cell gene expression Flex kit according to standard instructions. Libraries were sequenced on the NovaSeq6000 (Illumina) platform. Cell Ranger v.3.1.0 (10x Genomics) was used to align the sequencing reads to the GRCm38 mouse reference genome and generate count tables of unique molecular identifiers (UMIs) for each gene per cell.

#### Quality control and processing of single-cell RNA sequencing data

The R package Seurat v.4.1.1 was used to calculate the quality control metrics. Cells were removed from the analysis if a) there were fewer than 500 or greater than 3000 unique genes, b) there were more than 20,000 RNA molecules, or C) more than 8% of the reads mapped to mitochondrial genes. The data were normalized for sequencing depth, scaled to 10,000 UMIs per cell and log-transformed using the Seurat normalization data function. The Seurat FindVariableFeatures function was used to define 2000 highly variable genes that were used as input for the principal component analysis. Normalized data were scaled, and elbow plots were generated to determine how many principal components to include in the analysis. This corresponded to roughly the first 15 principal components. Uniform manifold approximation and projection (UMAP) embeddings were calculated using these principal components as input, and cells were clustered using the FindClusters function. We further filtered cells from clusters with fewer than 50 cells and repeated the above processing steps to identify our clusters of interest. *Single-cell RNA sequencing cluster identification and expression analysis* Clusters were annotated based on comparisons to neutrophil populations identified in a previous study of murine neutrophils (52). Briefly, gene signatures from the populations identified in *Xie* et al. were scored in each of the 5 detected clusters in our study using Seurat’s AddModuleScore function. Mature clusters corresponded to those scoring highly for G4-G5C, while immature populations scored highly for G1. Markers for each labeled population were identified using the quickMarkers function from the SoupX R package and filtered for those expressed in at least 65% of cells (52). These markers and other specific neutrophil markers were visualized using Seurat’s DoHeatmap and FeaturePlot functions.

### Adenosine quantification assay

Neutrophils were treated with fatty acids as indicated in the figure legends for 24 hours, after which the cells were centrifuged to obtain the supernatant. The supernatant was immediately assessed for targeted adenosine quantification using an adenosine assay kit (CELL BIOLABS, INC.) according to the manufacturer’s instructions.

### Real-time quantitative PCR

For neutrophil gene expression, one milligram of total RNA was extracted from neutrophils using a Quick RNA MicroPrep Kit (Zymo Research) according to the manufacturer’s instructions and reverse transcribed to cDNA using the SuperScriptTM III First-Strand Synthesis System (Invitrogen). Three technical replicates of real-time PCRs were performed for each sample. cDNA gene-specific primers and SYBR Green master mix (Applied Biosystems) were combined and then run on a QuantStudio 5 Real-Time PCR System (Applied Biosystems). The relative expression of the genes was calculated using the ddCt method with Actb or GAPDH as the housekeeping gene mRNA.

For tumor gene expression, MC38 cancer cells were subcutaneously injected into the flanks of C57BL/6 mice. Mice were sacrificed at either 14 days post injection (early time point) or 28 days post injection (late time point). Tumors were rapidly dissected. 1 microgram of total RNA was reverse transcribed to cDNA using the SuperScriptTM III First-Strand Synthesis System (Invitrogen). Three technical replicates of real-time PCRs were performed for each sample. cDNA gene-specific primers and SYBR Green master mix (Applied Biosystems) were combined and then run on a QuantStudio 5 Real-Time PCR System (Applied Biosystems). The relative expression of the genes was calculated using the ddCt method with Actb as the housekeeping gene mRNA.

### ELISA

Cytokine levels were quantified in supernatants from T cells after coculture with neutrophils from either tumor-bearing mice or mice induced with fatty acids. Mouse TNF-α, IFN-γ, IL-2, IL-17, IL-4, IL-10 and TGF-β assays were performed using paired antibody ELISA kits (R&D Systems, Minneapolis, MN) according to the manufacturer’s protocol.

### Data presentation and statistical analysis

The results are expressed as the mean ± standard deviation, unless otherwise stated. The significance of differences between two groups was tested using a 2-tailed, unpaired *t* test. Significant differences among multiple groups were tested using 1-way or 2-way ANOVA followed by Tukey’s post hoc test for multiple comparisons. A P value less than 0.05 was considered to indicate statistical significance. The statistical significance of the differences is described in the figure legends as *P < 0.05, **P < 0.01, ***P < 0.001 and *** P<0.0001.

## Results

### Neutrophils in the late-stage TME are immunosuppressive

Many solid tumors commonly exhibit a highly immunosuppressive TME, which causes dysregulation of the antitumor immune response and contributes to the ineffectiveness of ICB therapies. To determine whether the timing of anti-PD-L1 therapy influences therapeutic outcome, we administered anti-PD-L1 monoclonal antibody (mAb) on either day 4 (early) or day 14 (late) following implantation of syngeneic tumor lines (Figure 1A). Early anti-PD-L1 blockade reduced tumor volume in MC38- and Yumm5.2-bearing mice, while late anti-PD-L1 blockade had no impact on tumor growth (Figure 1B-C and S1A-B). Moreover, CD8⁺ T cells isolated from tumors of mice treated early exhibited enhanced effector function compared to those from late-treated tumors (Figure 1D). These results suggest that temporal changes in the TME may drive increasing immunosuppression and resistance to ICB.

**Figure 1.**
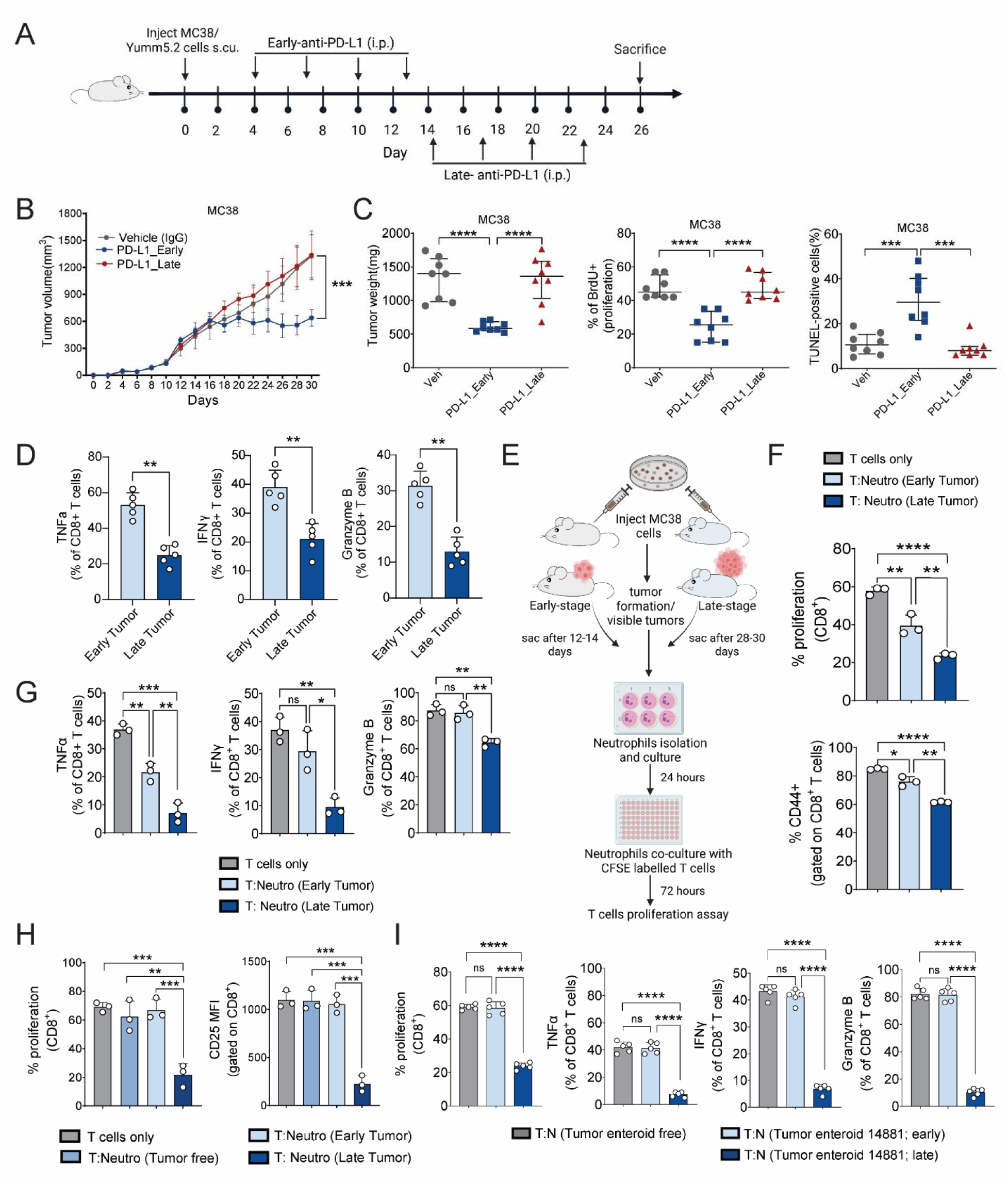
Late-stage neutrophils in the TME display immunosuppressive properties. (A) Schematic of the early and late anti-PD-L1 treatment regimens used in MC38 tumor-bearing mice. (A-E) WT C57BL/6 mice were injected subcutaneously with 2 X 10^6^ million MC38 tumor cells on both flanks on day 0, and after 3 days, anti-PD-L1 or IgG1 isotype Abs were administered intraperitoneally at a dose of 100 μg per mouse. A total of 4 doses were given every 3 days until the 13^th^ day for early treatment. The late anti-PD-L1 treatment started on the 14th day and lasted until the 23rd day for a total of 4 doses. (B) Tumor volume measured in MC38 tumor-bearing mice (n=8) (C) Tumor weight, proliferation and apoptosis were measured in MC38 tumor-bearing mice (n=8). (D) The effector functions (percentage of TNFα-, IFNγ- and Granzyme B-positive cells) on CD8+ cells harvested from MC38 subcutaneous tumors (n=5, biological replicates) were measured via flow cytometry. (E) Schematic showing neutrophil isolation from early-stage (day 14) and late-stage (day 30) MC38 tumor-bearing mice and subsequent coculture with CD3+ T cells at a ratio of 1:2 (neutrophil:T-cell). (F-G) Effect of neutrophils isolated from tumors of MC38 tumor-bearing mice (n=3, biological replicates) on *in vitro* T-cell functions (F) T-cell proliferation (percentage of CFSE dim/negative cells) and activation (percentage of CD44-positive cells) and (G) effector functions (percentage of TNFα-, IFNγ- and Granzyme B-positive cells) measured in CD8+ T cells via flow cytometry. (H) Effect of neutrophils isolated from the bone marrow of MC38 tumor-bearing mice (n=3, biological replicates) on *in vitro* T-cell proliferation (percentage of CFSE dim/negative cells) and activation (mean fluorescence intensity of the CD25 surface marker) measured in CD3+CD8+ cells through flow cytometry. (I) Effect of neutrophils isolated from established human tumor enteroids (n=5, biological replicates) on *in vitro* T-cell proliferation (percentage of CFSE dim/negative cells) and effector functions (percentage of TNFα-, IFNγ- and Granzyme B-positive cells) measured in CD3+CD8+ cells through flow cytometry. The data represent the mean ± s.d. *p < 0.05, ** p < 0.01, *** p < 0.001, **** p <0.0001.

Recent studies have identified neutrophil heterogeneity as a critical determinant of immunotherapy success (16,17), prompting us to investigate whether changes in neutrophil phenotype over time contribute to therapeutic resistance. Neutrophils isolated from tumor tissue of early- or late-stage MC38 tumor-bearing mice were cocultured with T cells, and their impact on T-cell proliferation, activation and effector function was evaluated (Figure 1E). Neutrophils from late-stage tumor-bearing mice, compared to early-stage tumors, more potently suppressed CD4+ and CD8+ T cell proliferation and activation (Figure 1F, Figure S1C for gating strategy, & S1F). Furthermore, the frequency of TNF-α, IFN-γ, and granzyme B positive CD8 T cells decreased in cells cultured with late-stage tumor neutrophils compared to those cultured with early-stage tumor neutrophils (Figure 1G). We also found that neutrophils from late-stage tumors increased the exhaustion molecules PD1 and TIM3 on CD8+ T cells, thereby promoting a dysfunctional state in cytotoxic CD8 T-cells (Figure S1G). Gene expression analysis further revealed that neutrophils from late-stage tumors displayed an immunosuppressive transcriptional profile, with reduced neutrophil elastase and myeloperoxidase and elevated arginase, iNOS, IL-10, and TGF-β expression (Figure S1D–E).

To assess whether this phenotype was restricted to the tumor or reflected systemic changes, we examined neutrophils from the bone marrow (BM). Strikingly, BM neutrophils from late-stage tumor-bearing mice also suppressed CD4⁺ and CD8⁺ T-cell proliferation and CD25 expression, whereas neutrophils from tumor-free or early-stage mice did not (Figure 1H and S1H), suggesting that late-stage tumors systemically reprogram neutrophils. This immunosuppressive phenotype also extended to tumor-associated macrophages from the late-stage TME, which similarly suppressed T-cell proliferation and activation (Figure S1I). These findings demonstrate that progressive immunosuppressive reprogramming of myeloid cells, particularly neutrophils and macrophages, occurs even within CMS1-like tumors, and is likely a general feature of tumor evolution that contributes to immunotherapy resistance.

Given that advanced colorectal tumors are often associated with increased stromal remodeling and may adopt a CMS4-like transcriptional program, we assessed whether late-stage tumors in our models exhibited features consistent with this phenotype. Specifically, we analyzed the expression of epithelial and mesenchymal markers commonly associated with epithelial-to-mesenchymal transition (EMT), a hallmark of CMS4 tumors. While late-stage tumors showed reduced expression of epithelial markers (CDH1, KRT20, EpCAM) compared to early-stage tumors, there was no corresponding increase in mesenchymal markers (SNAI1, VIM, TWIST1) (Figure S1J-K). These data indicate that late-stage tumors in our models do not undergo a transcriptional shift toward a CMS4-like state and suggest that the enhanced immunosuppressive phenotype is not driven by a mesenchymal transition. Instead, our findings point to functional reprogramming of the myeloid compartment, particularly neutrophils, as a central driver of immune suppression in the late-stage tumor microenvironment.

To determine whether the observed neutrophil reprogramming was simply a reflection of the immune-responsive nature of syngeneic lines, we extended our analysis to a patient-derived tumor enteroid model. Neutrophils isolated from late-stage tumor enteroid displayed potent immunosuppressive activity, significantly reducing CD8⁺ T-cell proliferation and effector function compared to neutrophils isolated from mice with early-stage enteroids (Figure 1I).

### Nutrients in the TME reprogram naïve neutrophils to become immunosuppressive

We investigated whether nutrients within the TME contribute to neutrophil reprogramming. We treated neutrophils with tumor interstitial fluid (TIF) or serum from MC38 colon cancer and Yumm5.2 melanoma tumor-bearing mice (Figure 2A). TIF-induced neutrophils significantly reduced CD4+ and CD8+ T-cell proliferation (Figure 2B), while serum-educated neutrophils also had a significant but more moderate impact (Figure S2A). Additionally, T cell effector function was compromised in the presence of TIF-educated neutrophils, as indicated by decreased IFNγ and TNFα levels (Figure 2C). To determine whether these effects extend to human disease, we examined TIF from human colorectal adenoma and adenocarcinoma tissues. Neutrophils educated with human adenocarcinoma TIF showed enhanced immunosuppressive activity, significantly reducing CD8+ T-cell proliferation and effector functions compared to those treated with adenoma TIF or controls (Figure 2D). These findings highlight a conserved role for TIF in driving neutrophil-mediated immunosuppression across murine models and human colorectal cancer.

**Figure 2.**
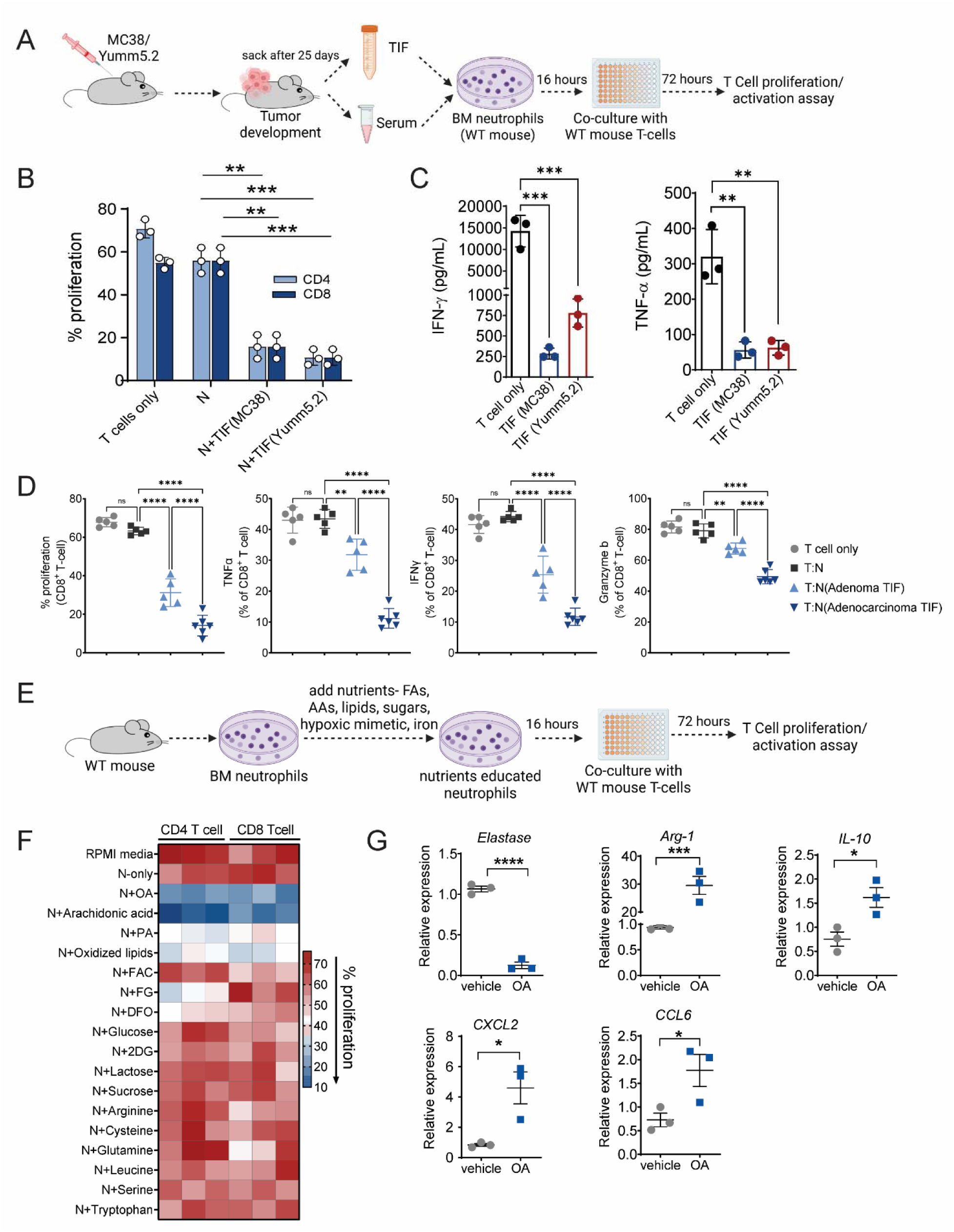
Naïve neutrophils are programmed by nutrients in the TME. (A) Schematic showing the isolation of tumor interstitial fluid (TIF) and serum from tumors of MC38-bearing C57BL/6 mice and the subsequent coculture of TIF-educated neutrophils with CD3+ T cells from WT C57BL/6 mice. (B-C) Effect of TIF-educated neutrophils (n=3, biological replicates) on *in vitro* T-cell functions, (B) CD4+ and CD8+ T-cell proliferation (percentage of CFSE dim/negative cells), and (C) IFN-γ and TNF-α secretion by CD3+ T cells when cocultured with TIF-educated neutrophils, as measured by ELISA. (D) Effect of patient TIF-educated human neutrophils (n=5, biological replicates) on *in vitro* T-cell proliferation (percentage of CFSE dim/negative cells) and effector functions (percentage of TNFα-, IFNγ- and Granzyme B-positive cells) measured in CD8+ cells when cocultured with TIF-educated neutrophils, measured via flow cytometry. (E) Schematic showing the isolation of bone marrow-derived neutrophils from WT C57BL/6 mice (n=3, biological replicates) and their treatment with various nutrients following subsequent coculture of nutrient-educated neutrophils with CD3+ T cells from WT C57BL/6 mice. (F) Heatmap showing the effect of nutrient-educated neutrophils on CD4+ and CD8+ T-cell proliferation. (G) Real-time qPCR validation of activation and immunosuppressive genes in cultured bone marrow-derived neutrophils (n=3, biological replicates) induced with 100 μM oleic acid (OA) or vehicle for 16 hours. The data are presented as the means ± s.d. *p < 0.05, ** p < 0.01, *** p < 0.001, **** p <0.0001.

Our data support the concept that neutrophils are metabolically plastic and can be functionally reprogrammed by local cues in the TME. The suppressive effects of MC38 and Yumm5.2 TIFs persist after heat inactivation (Fig S2B-C), suggesting that heat-stable components such as metabolites or lipid species, rather than proteins, mediate this effect. Thus, we conducted a targeted nutrient screen focusing on metabolic regulators that might augment neutrophil immunosuppression (Figure 2E). Fatty acids such as oleic acid (OA), arachidonic acid (AA), palmitic acid (PA), and oxidized lipids significantly increased neutrophil immunosuppressive activity, as evidenced by the decreased proliferative capacity of CD4+ and CD8+ T cells (Figure 2F). This was corroborated by changes in gene expression, as indicated by decreased elastase and increased arginase-1 and IL-10 levels in neutrophils treated with OA, as well as elevated chemokines CXCL2 and CCL6 linked to neutrophil migration into the TME (Figure 2G). Collectively, these results underscore the role of fatty acid metabolism in driving neutrophil immunosuppressive programming within the tumor microenvironment.

### In the late-stage TME, adenosine metabolism is modulated by fatty acids

To explore neutrophil heterogeneity in advanced tumors, we conducted single-cell RNA sequencing (scRNA-seq) on neutrophils from five late-stage tumor-bearing mice, pooled to minimize inter-sample variation. After quality control, we obtained 7,654 high-quality cells expressing a median of 1,467 genes per cell, detecting 14,813 unique mouse genes. Unbiased clustering identified five distinct populations (Figure S3A). To annotate these, we applied transcriptional signatures from Xie et al. (52), Based on these signatures, clusters 0/1 corresponded to mature neutrophils and clusters 2/3/4 to progenitor-like subsets (Figure S3B–C). We next examined marker expression, including CD39 (Entpd1) and CD73 (Nt5e) ectoenzymes critical for adenosine production. CD39 was enriched in a distinct subset, while CD73 showed lower but broader expression (Figure S3D–F). Both mature populations expressed canonical neutrophil markers Ly6G and Itgam (CD11b). CD39 and CD73 together convert extracellular ATP to immunosuppressive adenosine, which dampens immune responses via purinergic signaling.

To assess conservation in human disease, we analyzed scRNA-seq data from colorectal and pancreatic cancer patients. In colorectal tumors, specific myeloid populations were extracted from a colorectal carcinoma single cell dataset(53). UMAP of 5 major myeloid populations of interest from 63 patient tumor samples were identified (Figure S3G). CD39 was enriched in CD11c⁺ dendritic cells, activated neutrophils, and granulocytes, while CD73 remained low across immune populations (Figure S3G–I). High CD39 expression correlated with immunosuppressive markers including TGFB1 and PTGS2 (Figure S3J). Similar patterns were seen in pancreatic cancer, where granulocytes and activated neutrophils again showed elevated CD39 (Figure S3K–N)(54). These data support a conserved role for CD39 in shaping the immunosuppressive myeloid compartment in cancer.

We validated these findings using flow cytometry, confirming an upregulation of CD39 and CD73 in tumor-associated neutrophils (Figure 3A) from late-stage tumor-bearing mice compared to those from early-stage tumors. Similar patterns were observed in BM neutrophils (Figure S3O). Together, these data suggest that late-stage neutrophils adopt a transcriptional and functional program crucial for adenosine generation.

**Figure 3.**
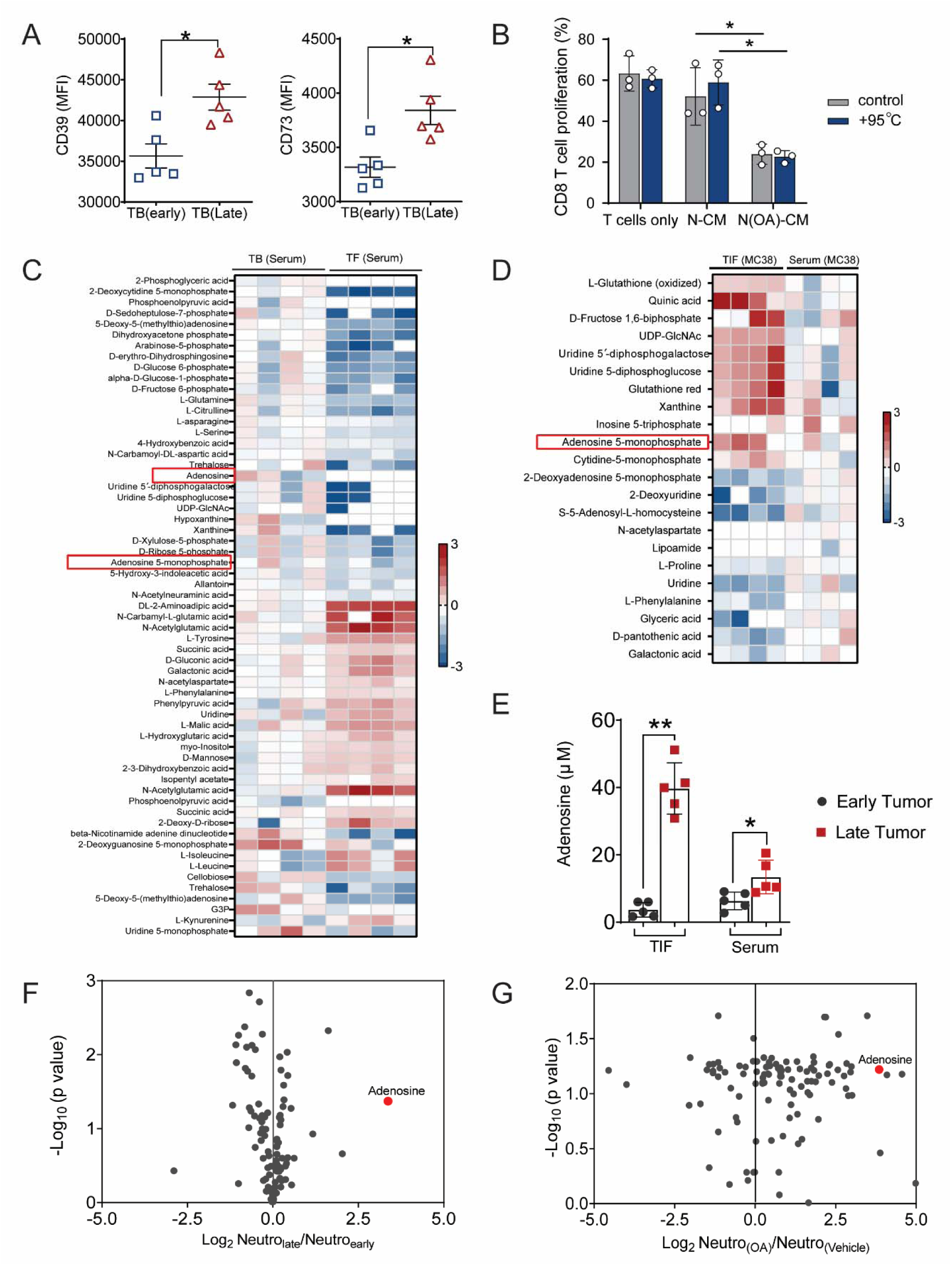
Neutrophils or TIFs from the late-stage tumor microenvironment show enrichment of the adenosine pathway regulated via oleic acid. (A) Mean fluorescence intensity (MFI) of CD39 and CD73 expression measured via flow cytometry in tumor neutrophils (7AAD-CD45+CD11b+Ly6G+) (n=5, biological replicates) isolated from MC38 tumor-bearing early-stage or late-stage mice. (B) Assessment of CD8+ T-cell proliferation via flow cytometry after treatment with OA-induced neutrophil control or heat-inactivated (95°) conditioned media at a ratio of 3:1 (conditioned media from neutrophils: regular T-cell media). (C-D) Heatmap plots showing significant metabolite abundance levels analyzed through metabolomics for comparison between (C) MC38 tumor-bearing mouse serum and tumor-free mouse serum and (D) TIF and serum from MC38 tumor-bearing mice (n=4, biological replicates in Figure C or D). (E) Quantification of adenosine measured through a fluorescence-based assay in TIFs and serum from early- or late-stage tumor-bearing mice. (F-G) Representative volcano plots showing significant metabolites in (F) late-stage neutrophils with respect to early-stage tumor neutrophils from MC38 tumor-bearing mice (n=4). (G) OA-induced bone marrow neutrophils with respect to vehicle-treated neutrophils from WT C57BL/6 mice (n=4). The data represent the mean ± s.d. *p < 0.05, ** p < 0.01.

To identify suppressive factors secreted by these neutrophils, we analyzed conditioned media from OA-treated neutrophils. Heat inactivation failed to abrogate T cell suppression (Figure 3B, S3P), implicating metabolites rather than proteins. Since CD39 and CD73 are rate-limiting for adenosine production, we next quantified adenosine via metabolomics. Serum from tumor-bearing mice exhibited elevated adenosine and AMP compared to tumor-free controls (Figure 3C), with TIF containing even higher AMP levels than serum (Figure 3D). Both TIF and serum from late-stage tumors showed greater adenosine than those from early-stage tumors (Figure 3E). Additionally, conditioned media from neutrophils isolated from late-stage tumors, as well as OA-treated neutrophils, contained significantly more adenosine than their respective controls (Figure 3F–G). Together, these findings demonstrate that fatty acid exposure and tumor progression drive neutrophil reprogramming toward an adenosine-producing, immunosuppressive phenotype in the TME.

### Fatty acids regulate the neutrophil-adenosine axis in the late-stage tumor microenvironment

Our nutrient screen revealed that fatty acids reprogram naïve neutrophils into immunosuppressive cells, which aligns with findings from other studies implicating fatty acids, such as arachidonic acid, in the suppressive activity of myeloid-derived suppressor cells (MDSCs) (55). We analyzed the local lipidome of TIFs from MC38 tumor-bearing hosts using comprehensive lipidomics (Table S1-S2). To define the local lipid landscape, we performed lipidomics on TIFs from early- and late-stage MC38 tumors and normalized these profiles to serum from the same tumor-bearing mice (Tables S3). Given that serum from late-stage tumors also exhibited immunosuppressive properties, albeit less pronounced than in TIFs, we performed comparative lipidomic profiling. We independently compiled lists of lipid species significantly altered in serum and in TIFs between early- and late-stage tumors and then intersected these datasets to identify lipid species commonly dysregulated in both compartments during tumor progression (Table S4). Together, the data demonstrates that extensive lipid reprogramming locally and systemically. To further investigate the spectrum of lipids and the underlying mechanisms by which fatty acids influence adenosine production in neutrophils, we performed a lipid screen. Neutrophils were supplemented with 80 different fatty acids, and adenosine levels were measured in the conditioned media. This analysis revealed that linoleic acid (LA), alpha-linolenic acid (ALA), arachidonic acid (AA), oleic acid (OA), and their esterified derivatives were the most effective at increasing adenosine production in neutrophils (Figure 4A). These fatty acids also upregulated CD39 and CD73 expression in neutrophils (Figure 4B) and impaired CD4⁺ and CD8⁺ T-cell proliferation and effector functions (Figure 4C–D), without inducing significant neutrophil death (Figure S4B). While neutrophils and macrophages both exhibited increased adenosine production following fatty acid exposure (Figure S4A), macrophages also adopted a suppressive phenotype capable of inhibiting T-cell responses (Figure S4C), suggesting broader myeloid immunosuppression within the late-stage TME.

**Figure 4.**
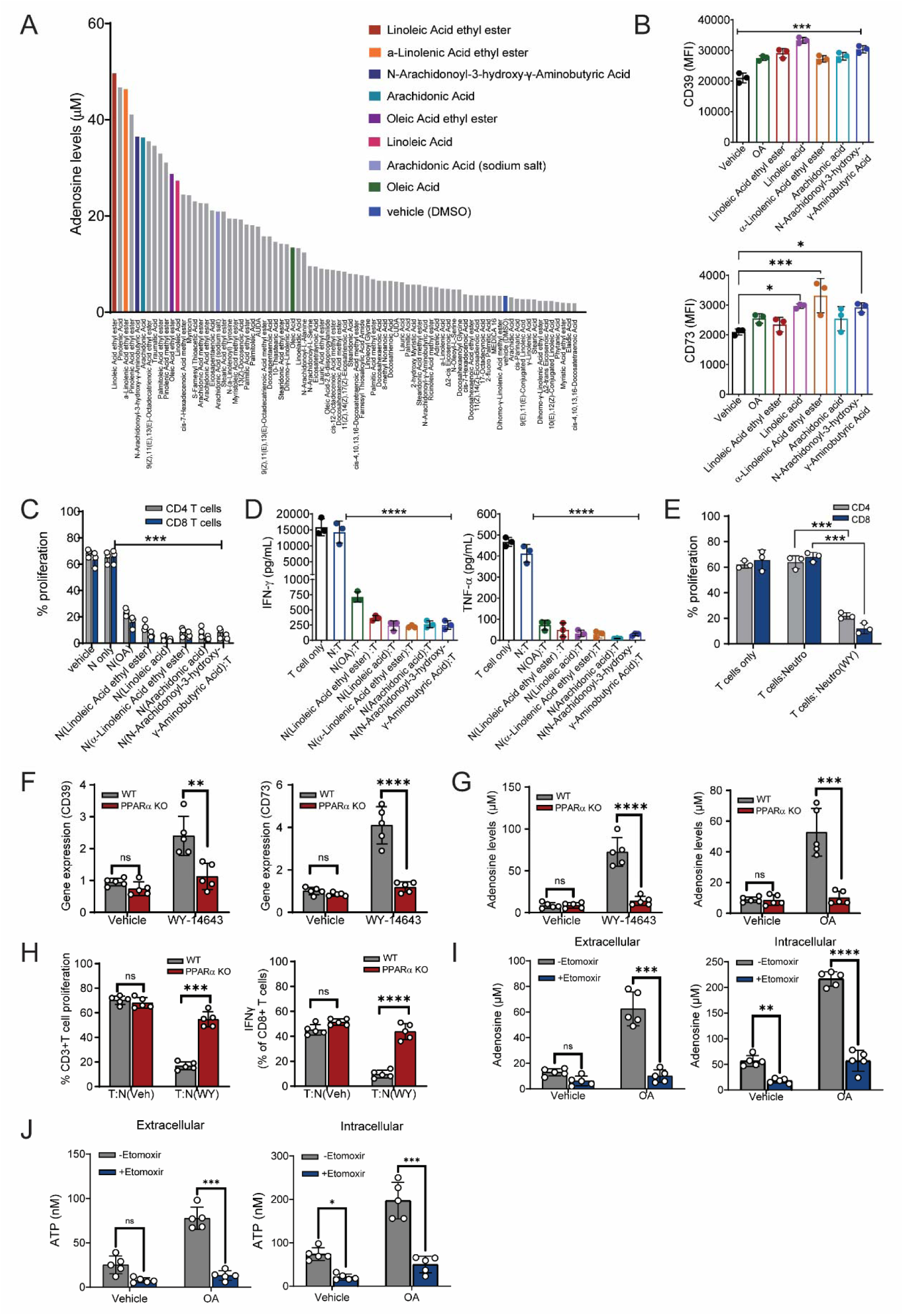
Neutrophil adenosine secretion is regulated through unsaturated fatty acids via PPAR**α**. (A) Quantification of adenosine levels measured through a fluorescence-based assay in conditioned medium from bone marrow-derived neutrophils of WT C57BL/6 mice (n=3 independent mice) treated with 50 μM of the 80 different indicated fatty acids for 16 hours. Each bar represents an average value of three triplicate measurements of adenosine from conditioned media. (B) Flow cytometric measurements of the levels of the adenosine synthesis enzymes CD39 and CD73 in bone marrow-derived neutrophils (7AAD-CD45+CD11b+Ly6G+) (n=3 independent mice) treated with 50 μM of the indicated fatty acids for 16 hours. (C-D) Effect of the indicated fatty acid-educated neutrophils (n=3, biological replicates) on *in vitro* T-cell function, (C) CD4+ and CD8+ T-cell proliferation (percentage of CFSE dim/negative cells), and (D) TNF-α and IFN-γ secretion by CD3+ T cells when cocultured with fatty acid-educated neutrophils, as measured by ELISA. (E) Effect of the WY-14643 treated neutrophils (n=3, biological replicates) on *in vitro* T-cell proliferation (percentage of CFSE dim/negative cells). (F-H) Treatment of WY-14643 (100μM) in WT and PPARα KO mice. (F) Measurements of CD39 and CD73 mRNA levels by q-PCR in neutrophils (n=5, biological replicates), (G) Adenosine levels, and (H) CD3+ T cell proliferation and IFN-γ secretion. (I) Quantification of extracellular and intracellular adenosine levels in vehicle- or OA-conditioned neutrophils after treatment of etomoxir. (J) Quantification of extracellular and intracellular ATP levels upon vehicle- or OA-conditioning in neutrophils. The data are presented as the means ± s.d. *p < 0.05, ** p < 0.01, *** p < 0.001, **** p <0.0001.

Fatty acids can activate peroxisome proliferator-activated receptors (PPARs), a family of ligand-activated transcription factors (56,57). Among these, OA selectively activates PPARα signaling (58). Consistent with this, the PPARα agonist WY-14643 induced potent immunosuppressive activity in neutrophils, reducing both CD4⁺ and CD8⁺ T-cell proliferation (Figure 4E) and activation (Figure S4D). WY-14643 upregulated CD39 and CD73 expression and increased adenosine production in wild-type but not PPARα-deficient neutrophils (Figure 4F–G), confirming a PPARα-dependent mechanism(57–60). Furthermore, suppression of T cell proliferation and IFNγ production was abolished in PPARα knockout mice (Figure 4H), highlighting the requirement of this pathway for fatty acid-driven immunosuppressive programming.

Given that ATP generated via β-oxidation serves as a substrate for adenosine, we inhibited β-oxidation with etomoxir and observed reduced intra- and extracellular adenosine levels in fatty acid-treated neutrophils (Figure 4I, S4E). OA also significantly elevated intracellular and extracellular ATP in neutrophils (Figure 4J), supporting the link between fatty acid metabolism and adenosine generation. Notably, OA did not induce neutrophil extracellular trap (NET) formation (Figure S4F), indicating that ATP increases were not NET-dependent.

Together, these findings define a mechanism by which fatty acids in the late-stage TME reprogram neutrophils via PPARα-dependent metabolic signaling to produce adenosine and suppress antitumor immunity.

### Targeting the adenosine axis reverses neutrophil-mediated T-cell suppression

Having established that fatty acids drive neutrophil-mediated adenosine production through PPARα, we next assessed whether pharmacologic inhibition of the adenosine signaling axis could restore T-cell function. To disrupt this pathway at multiple nodes, we employed two agents: AB680, a potent CD73 inhibitor that blocks AMP-to-adenosine conversion, and AB928, an orally available dual A2aR/A2bR antagonist currently in clinical trials (Figure 5A). We first confirmed that AB680 effectively reduced extracellular adenosine levels in OA-treated neutrophils without altering intracellular adenosine concentrations (Figure S5A). To evaluate whether pharmacologic blockade of the adenosine pathway could restore T cell function, neutrophils were pretreated with AB680, or T cells were pretreated with AB928.

**Figure 5.**
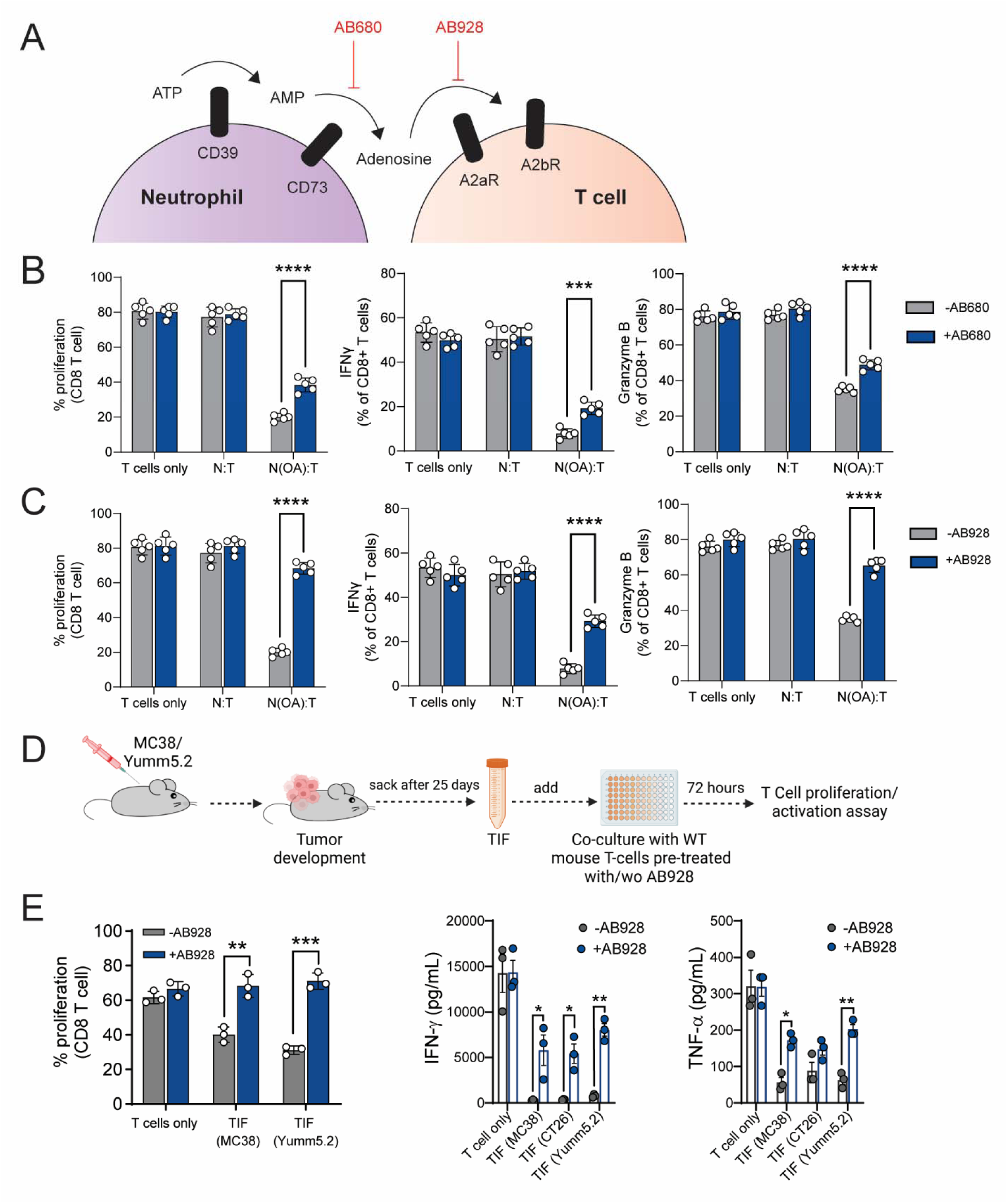
Adenosine pathway inhibition via AB680 or AB928 rescues the suppressive effects of neutrophils on T cells. (A) Schematic showing the targets of AB680 and AB928in adenosine pathway. (B-C) Effect of the indicated fatty acid-educated neutrophils (n=5, biological replicates) on *in vitro* (B) AB680-pretreated or (C) AB928-pretreated T-cell functions such as proliferation (percentage of CFSE dim/negative cells) and effector functions (IFN-γ and Granzyme B secretion) on CD8+ T cells when cocultured with fatty acid-educated neutrophils, measured by flow cytometry. (D) Schematic showing the isolation of TIFs from tumors of MC38- or Yumm5.2- mice (n=3, independent mice for each cell line) and treatment of AB928-pretreated CD3+ T cells from WT C57BL/6 mice with TIFs. (E) Effect of TIF from the indicated tumor cells on *in vitro* AB928-pretreated T-cells. Cell proliferation (percentage of CFSE dim/negative cells) and effector functions (IFN-γ and TNF-α secretion) on CD8+ T cells was measured by Flow cytometry and ELISA. The data are presented as the means ± s.d. *p < 0.05, ** p < 0.01, *** p < 0.001, **** p <0.0001.

Immunosuppressive neutrophils were generated by treatment with OA, LA, AA, or their derivatives, followed by co-culture with T cells. While AB680 partially rescued T cell proliferation, AB928 more effectively reversed neutrophil-mediated suppression (Figure 5B–C). Furthermore, AB928 significantly enhanced T cell effector responses, as evidenced by increased IFNγ and granzyme B production. Although TNFα levels were also elevated, but no change in exhaustion markers following AB928 treatment, this effect varied depending on the fatty acid stimulus (Figure S5B-C).

Subsequently, we investigated whether adenosine is the major immunosuppressive metabolite in the TME. T cells were pretreated with AB928 or vehicle and cultured in the presence of TIFs isolated from MC38 and Yumm5.2 tumors (Figure 5D). TIFs suppressed CD4⁺ and CD8⁺ T-cell proliferation, an effect largely reversed by AB928 (Figure 5E, S5D). AB928 also enhanced effector cytokine production, including IFNγ and TNFα (Figure 5E). Together, these findings highlight the central role of adenosine in mediating immune suppression within the TME and demonstrate that dual A_2a_R/A_2b_R receptors blockade with AB928 offers superior immunomodulatory potential over upstream inhibition strategies.

### Fatty acid-induced neutrophils inhibit T-cell killing via adenosine

To assess how fatty acid–conditioned neutrophils influence CD8⁺ T cell–mediated tumor killing, we utilized the OT-I transgenic mouse model, in which CD8⁺ T cells express an ovalbumin (OVA)-specific TCR. Splenic OT-I CD8⁺ T cells were isolated and cultured for 3 days in conditioned media from neutrophils treated with fatty acids, then co-cultured with OVA-expressing MC38 colon carcinoma cells. T cell cytotoxicity was assessed by SYTOX Green staining and flow cytometry (Figure 6A–C). As expected, CD8⁺ T cells alone induced robust cancer cell death, and naïve neutrophils had no effect. In contrast, CD8⁺ T cells exposed to neutrophils preconditioned with fatty acids or TIFs from MC38, Yumm5.2, or CT26 tumors showed significantly impaired tumor cell killing (Figure 6B–C). Strikingly, pretreatment with the A2aR/A2bR antagonist AB928 restored cytotoxic activity, rescuing T cell–mediated killing of MC38-OVA cells.

**Figure 6.**
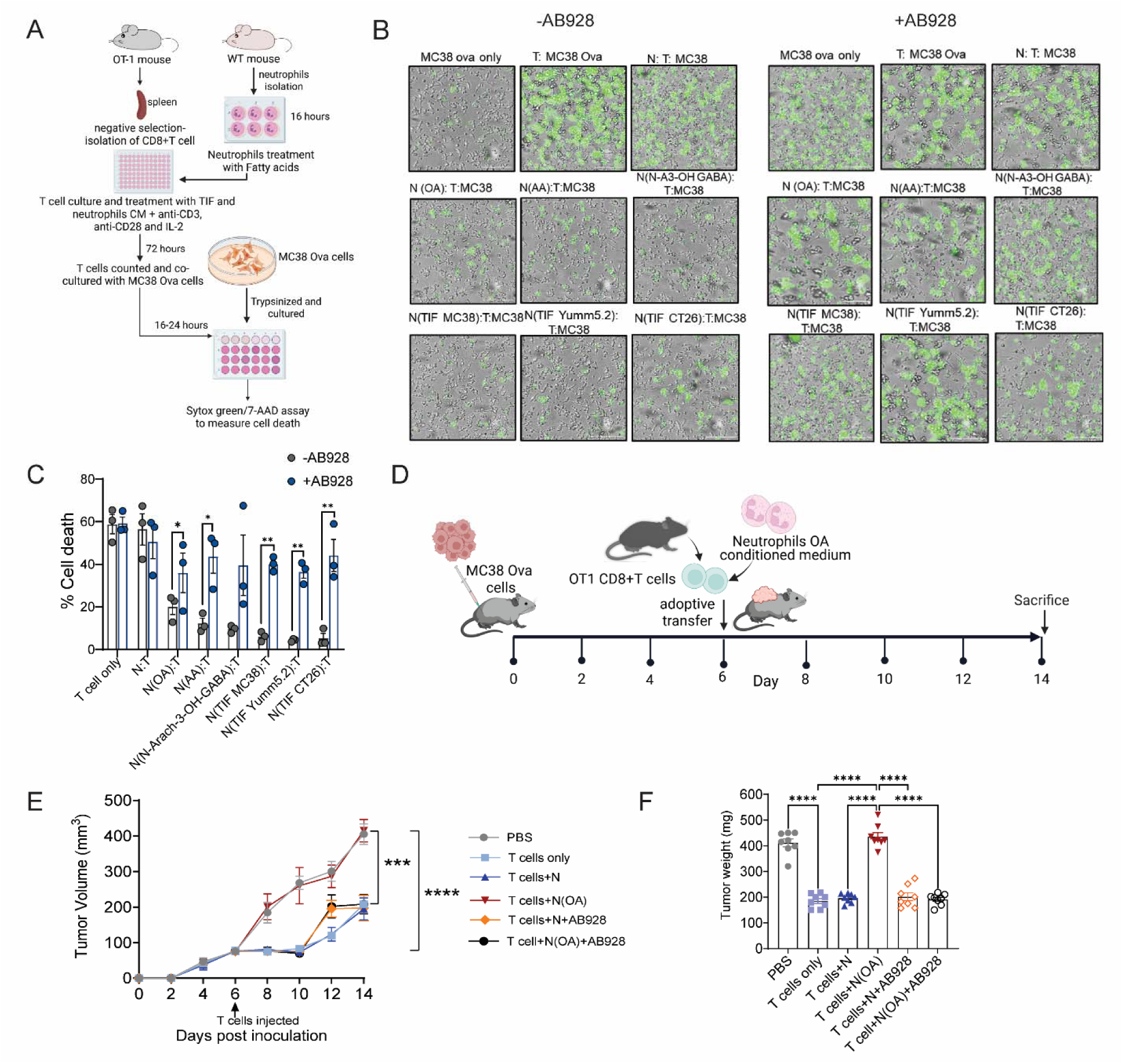
Fatty acid-induced neutrophils reduce the cytotoxic T cell killing capacity, which is dependent on adenosine. (A) Schematic of the experiments shown in panels (B-C). CD8+ T cells were isolated from OT-1 mice (n=3 technical replicates pooled from 5 individual OT-1 mice), pretreated with or without AB928 and cultured with conditioned medium from fatty acid- or TIF-educated bone marrow-derived neutrophils (pooled from 5 independent mice) for 3 days. The activated/suppressed CD8+ T cells were counted and then cocultured with MC38-Ova-expressing cells for 16 hours at a ratio of 4:1 (CD8+ T:MC38-Ova). (B-C) Effect of CD8+ T cells induced with conditioned medium from fatty acid/TIF-educated neutrophils on the viability of MC38-Ova cells. (B) Fluorescence-based assay showing SYTOX green-positive cells in microscopy images indicating dead cells. (C) Cell death measurements via 7AAD staining using flow cytometry. The data in panel C are presented as the means ± s.d.s; unpaired Student’s t tests were performed. (D) Schematic illustrating the CD8+ T-cell adoptive transfer experiment in which CD8+ T cells were isolated from OT-1 mice (n= 10, pooled from individual OT-1 mice), pretreated or not pretreated with AB928 and cultured with conditioned medium from OA (100 μM)-educated bone marrow neutrophils (pooled from 6 independent mice) for 4 days. T cells were counted, and 1 million cells per mouse for each different condition were intravenously injected into MC38-OVA tumor-bearing mice (n=0.8 for each group). The data are presented as the means ± s.d.s; two-way ANOVA with Tukey’s multiple comparison test was used for comparison. (E-F) Tumor growth measurements over time for MC38-Ova tumors. (E) Tumor volume measured for 14 days. (F) End-point tumor weight. The data are presented as s.e.m. *p < 0.05, ** p < 0.01, *** p < 0.001, **** p <0.0001.

To validate these findings in vivo, we performed adoptive transfer experiments in MC38-OVA tumor-bearing mice. OVA-specific CD8⁺ T cells were activated ex vivo in the presence or absence of OA-conditioned neutrophil media, with or without AB928 (Figure 6D). Control mice exhibited progressive tumor growth (∼500 mm³; Figure 6E). Mice receiving CD8⁺ T cells alone showed the greatest tumor suppression, with reduced tumor volume and weight, increased apoptosis, and decreased proliferation (Figure 6E–F, S6A–B). CD8⁺ T cells conditioned with control neutrophil media were similarly effective, indicating that naïve neutrophils do not impair T cell function. In contrast, tumors were resistant to CD8⁺ T cells exposed to OA-treated neutrophil media, as shown by increased tumor burden, reduced apoptosis, and elevated proliferation (Figure 6E–F, S6A–B). Notably, AB928 pretreatment rescued the cytotoxic function of CD8⁺ T cells, reducing tumor growth and restoring apoptotic and antiproliferative activity (Figure 6E–F, S6A–B).

Together, these findings demonstrate that fatty acid–programmed neutrophils suppress CD8⁺ T cell cytotoxicity via adenosine signaling, and that A2aR/A2bR blockade can restore antitumor immunity. Targeting this pathway may offer a promising strategy to enhance T cell–mediated tumor clearance in the context of advanced disease.

### Immunosuppressive neutrophils drive tumor progression in an autochthonous mouse model of colon cancer

Syngeneic tumor models do not completely recapitulate the TME observed in CRC. To validate our findings in a more clinically relevant model that better recapitulates human CRC progression and ICB resistance, we utilized an autochthonous metastatic model of CRC mediated by the deletion of *Apc* and *tp53* and the activation of mutant *Kras (TripleMut)* utilizing a tamoxifen-inducible colon epithelial-specific CDX2-CreER^T2^ model (42,61). This autochthonous CRC model closely mimics human disease progression and is highly resistant to ICBs (42,62). *TripleMut* mice were sacked 5 days (early-stage) or 10 days (late-stage) after tamoxifen treatment. The effects of neutrophils from colon tissue on T-cell functions were assessed (Figure 7A).

**Figure 7.**
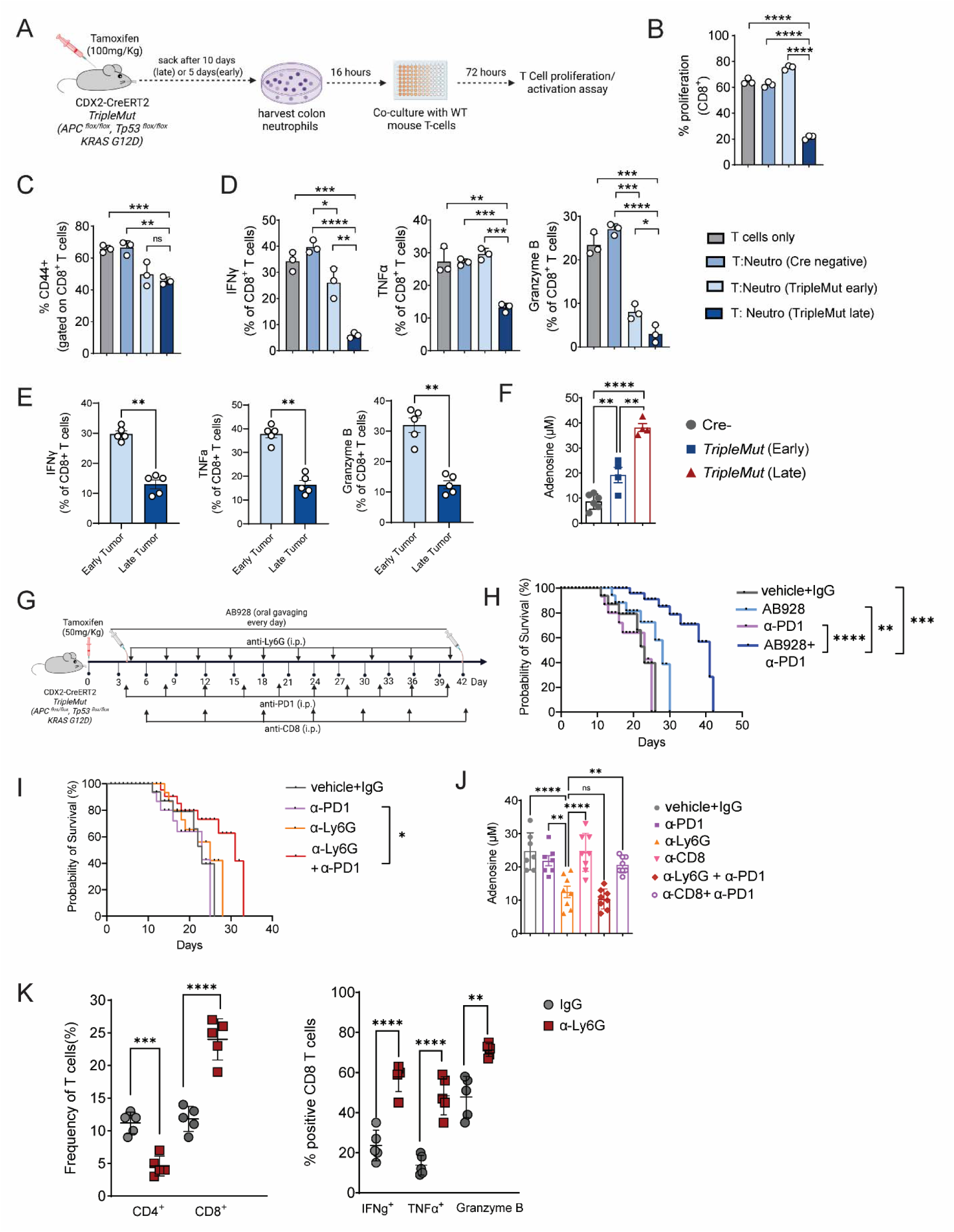
Blocking adenosine receptors increases the sensitivity of autochthonous CRC mice to ICB therapy. (A) Schematic illustrating colon neutrophil isolation from CDX2-ERT2Cre; *Apc^fl/fl^*; *Trp53^fl/fl^*; Kras^LSLG12D^ mice (*TripleMut*) induced with 100 mg/kg tamoxifen and subsequent coculture with CD3+ T cells from WT C57BL/6 mice. (B-D) Effect of TripleMut colon neutrophils (n=3, independent mice) on T-cell activity (B) proliferation (percentage of CFSE dim/negative cells), (C) activation (percentage of CD44-positive cells), and (D) effector functions (percentage of TNFα-, IFNγ- and Granzyme B-positive cells) measured in CD3+CD8+ cells via flow cytometry. (E) The effector functions (percentage of TNFα-, IFNγ- and Granzyme B-positive cells) measured on CD8+ T cells from early and late TripleMut tumors (n=5, biological replicates) via flow cytometry. (F) Adenosine quantification through a fluorescence-based assay in the serum of Cre-negative (n=6) *TripleMut early* mice sacked on the 4th day after tamoxifen injection (n=4) and *TripleMut late* mice sacked on the 9th day after tamoxifen injection (n=4). The data in panels B-F are presented as the means ± s.d.s; unpaired Student’s t tests were used. (G) Schematic illustrating the *in vivo* study of *TripleMut* mice. Mice were induced with a low dose of tamoxifen (50 mg/kg) for each group; after 3 days, they were orally gavaged twice a day with AB928 (100 mg/kg), treated intraperitoneally 3 days apart with anti-PD-1 (5 mg/kg) or the IgG2a isotype control, every 3^rd^ day with anti-Ly6G (100 μg/mouse) or 5 days apart with anti-CD8α (25 mg/kg) and monitored for survival (n=7-8 in each group). (H-I) Survival of *TripleMut* mice treated with an anti-PD-1 antibody or isotype control agent either alone or in combination with (H) AB928 (I) anti-Ly6G. Survival significance was calculated using a log-rank test. (J) Adenosine levels were measured in the serum of *TripleMut* mice treated with the indicated reagents. (K) T cells frequency and effector functions (percentage of IFNγ-, TNFα-, and Granzyme B-positive cells) measured in colon tissue of TripleMut mice following anti-Ly6G treatment. The data are presented as the means ± s.d. *p < 0.05, ** p < 0.01, *** p < 0.001, **** p <0.0001.

Neutrophils isolated from late-stage TripleMut colons significantly suppressed both proliferation and activation of CD4⁺ and CD8⁺ T cells compared to those from early-stage or Cre-negative mice (Figure 7B–C, S7A–B). Late-stage colon neutrophils also reduced the frequency of IFNγ⁺, TNFα⁺, and granzyme B⁺ CD8⁺ T cells (Figure 7D). In vivo analysis confirmed these findings, CD8⁺ T cells isolated from late-stage *TripleMut* tumors exhibited reduced effector function (Figure 7E), coinciding with elevated serum adenosine levels (Figure 7F). Additionally, colon neutrophils (Figure S7C) and macrophages (Figure S7D) from late-stage tumors upregulated the CD39 and CD73. Notably, immunosuppressive neutrophils also emerged in the bone marrow during late-stage disease (Figure S7E–F), reflecting systemic remodeling of the myeloid compartment.

Given that an immunosuppressive TME contributes to ICB resistance in CRC, and that fatty acid–driven adenosine production by neutrophils suppresses cytotoxic T cell activity, we hypothesized that targeting adenosine signaling could improve immunotherapy outcomes(60). To test this, *TripleMut* mice received tamoxifen (50 mg/kg), followed three days later by oral AB928 (A2aR/A2bR antagonist) and/or anti-PD1 treatment (Figure 7G). Neutrophils and CD8⁺ T cells were selectively depleted using anti–Ly6G or anti-CD8 antibodies, respectively (Figure 7G).

Anti-PD1 monotherapy failed to improve survival (Figure 7H, S7G), while AB928 alone modestly extended survival. Strikingly, combination treatment with AB928 and anti-PD1 led to a significant survival benefit (Figure 7H) and increased CD8⁺ T cell infiltration in blood and colon (Figure S7H). This effect was abolished by CD8⁺ T cell depletion (Figure S7G), confirming T cell dependence. Neutrophil depletion reduced circulating and colonic neutrophils (Figure S7I) and significantly improved survival when combined with anti-PD1 (Figure 7I, S7G). Correspondingly, serum adenosine levels were markedly reduced (Figure 7J), and frequencies of both CD4⁺ and CD8⁺ T cells with enhanced effector function were restored (Figure 7K). Together, these results demonstrate that neutrophils are a key source of extracellular adenosine in the CRC TME, contributing to immunosuppression and ICB resistance. Targeting neutrophil-driven adenosine signaling either pharmacologically or via cell depletion represents a promising strategy to improve immunotherapy efficacy in colorectal cancer.

## Discussion

The TME is a dynamic, metabolically active milieu that evolves over the course of tumor progression, ultimately leading to immunosuppressive surveillance. Our results reveal that the timing of ICB critically determines therapeutic efficacy. Anti–PD-L1 therapy is effective in early-stage tumors, but fails in advanced disease, where neutrophils acquire a potent immunosuppressive phenotype. Notably, this shift occurs in both the tumor and bone marrow compartment and extends to other myeloid cells, suggesting systemic reprogramming of the innate immune system. Mechanistically, this is attributed to a metabolic axis in which tumor-derived fatty acids reprogram neutrophils via PPARα-dependent signaling to suppress T cell function through adenosine production. This mechanism contributes to immunotherapy resistance in CRC and can be therapeutically targeted with adenosine receptor blockade.

Our nutrient screens, supported by lipidomic profiling and human TIF data, identify fatty acids particularly OA, LA, AA, and their derivatives as dominant drivers of neutrophil-mediated T cell suppression. These fatty acids induce adenosine secretion in a PPARα- and fatty acid oxidation-dependent manner, establishing a direct link between tumor lipid metabolism and innate immune suppression. Given that fatty acids are abundant in tumors and influenced by diet, host metabolism, and microbial contributions, it will be important to delineate how these factors shape the immunosuppressive neutrophil phenotype. Moreover, our findings reveal a temporal remodeling of the lipid composition within CRC TIFs, with distinct panels of lipids altered during tumor progression. While OA plays a key role, the identity and contribution of specific lipid species enriched in TIFs that drive neutrophil-mediated immunosuppression warrant further investigation. Moreover, serum from late-stage tumors induced some degree of T cell suppression via neutrophils. The observation that T cell suppression can be partially recapitulated by serum further supports the systemic nature of neutrophil reprogramming in cancer, which is further potentiated within the TME.

A recent study suggested that linoleic acid directly enhances CD8⁺ T cell memory and reverses their exhaustion phenotype (63). In contrast, another report showed that linoleic acid can inhibit CD4⁺ T cell functions in vitro (64). Our work demonstrates that linoleic acid and its derivatives reprogram neutrophils to produce adenosine, which suppresses both CD4⁺ and CD8⁺ T cell functions. Collectively, these findings highlight the complex role of fatty acids in modulating immune cell function and antitumor immunity. However, our data also suggest that lipids and more broadly, any nutrients within the TME must be evaluated in the context of heterogeneous cellular interactions. Metabolite-driven effects on CD8⁺ T cells may occur indirectly through other immune or stromal cell types, underscoring the need to consider multicellular metabolic crosstalk when assessing functional outcomes.

Adenosine is an immunosuppressive metabolite, and the upregulation of the adenosine pathway is a hallmark feature of many different types of cancers. Adenosine gene signatures are associated with worse outcomes in cancer patients (65–69). While adenosine is well known to inhibit T cell activation via A2aR/A2bR signaling, our data position neutrophils as a major and previously underappreciated source of extracellular adenosine in the TME. This function appears conserved in human colorectal and pancreatic tumors. The extent to which neutrophil-derived adenosine versus tumor- or stromal-derived adenosine dominates across cancers remains unclear. Moreover, how adenosine production by neutrophils is integrated with other immunosuppressive pathways, such as arginase, ROS, or PD-L1 expression requires further investigation.

Our findings also raise translational opportunities. The use of AB928, an A_2A_R antagonist, in phase 1/2 clinical trials for AB928-based combination therapies for many cancer types, including metastatic colorectal cancer, lung cancer, non-small cell lung cancer, prostate cancer, and other gastrointestinal malignancies, is under investigation (NCT04660812, NCT03846310, NCT04262856, NCT04381832, and NCT03720678). In our study, the A2aR/A2bR antagonist AB928 restored CD8⁺ T cell function and synergized with anti–PD-1 therapy in a resistant *TripleMut* CRC model. These results support ongoing clinical efforts to target adenosine signaling and suggest that biomarkers of lipid metabolism or neutrophil activation could stratify patients likely to benefit. However, the incomplete rescue seen with AB928 alone highlights the complexity of immune suppression in late-stage tumors. Future studies should evaluate combinatorial strategies that target upstream metabolic regulators (e.g., PPARα), neutrophil recruitment, or additional immunosuppressive metabolites.

In sum, we identify a fatty acid–driven, PPARα-mediated immunosuppressive program in neutrophils that promotes adenosine accumulation and T cell dysfunction. This axis is a critical driver of ICB resistance and represents a metabolically vulnerable node for therapeutic intervention. Targeting the neutrophil–adenosine axis may improve outcomes in CRC and other tumors characterized by a lipid-rich, immunosuppressive microenvironment.

## Study Limitations

Our study utilizes syngeneic tumor models, patient samples, patient-derived enteroids, and a genetically engineered autochthonous CRC model to explore neutrophil-mediated immunosuppression. However, these systems have inherent limitations in fully capturing the complexity and heterogeneity of human colorectal cancer. While comparing early and late tumor stages provides valuable insights, this approach may miss dynamic changes that occur throughout tumor progression. Furthermore, while we identified key lipids that drive immunosuppression, the specific contributions and interactions of the broader array of lipid species within the TME require further investigation.

## Supporting information

Figure S1. Myeloid cells in the tumor microenvironment display immunosuppressive gene signatures (related to Figure 1).

Figure S2. TIF and serum from advanced tumor-bearing mice induce immunosuppression through neutrophils (related to Figure 2).

Figure S3. Single-cell sequencing revealed heterogeneity in late-stage tumor neutrophils (related to Figure 3).

Figure S5. AB680 and AB928 abolishes the suppressive effects of fatty acid-induced neutrophils on T cells (related to Figure 5).

Figure S6. AB928 restored the T-cell killing capacity in vivo (related to Figure 6).

Figure S7. Myeloid cells display high expression of adenosine signatures in an advanced autochthonous CRC mouse model.

## Acknowledgments

A schematic diagram (Figures 1A, 1E, 2A, 2E, 5D, 6A, 6D, 7A and 7G) was created with <u>BioRender.com</u>. The authors thank Arcus Biosciences for providing AB680 and AB928 for the experiments. This work was funded by NIH grants R01CA148828, R01CA245546, R01DK095201, and UMCCC Core Grant P30CA046592 (Y.M.S.). R01AI152517 to C.A.P.

R01CA248160 to C.A.L. RS was supported by a Postdoctoral Fellowship (032650) from the American Physiological Society and a Research Fellow Award (1003279) from the Crohn’s and Colitis Foundation. H.N.B. was supported by an NCI Predoctoral Fellowship (5F30CA257292-02). S.S. was supported by a Crohn’s and Colitis Foundation Research Fellow Award (623914) and the American Heart Association Postdoctoral Fellowship (19POST34380588). W.H. was supported by a NIDDK F30 predoctoral grant (F30DK131851). MDG is supported by NIH (1R01CA276217) and VA (I01 BX005267) grants. ST and FJG were supported by the National Cancer Institute Intramural Research Program (ZIABC005562-36). CIKC was supported by JSPS Research Fellowship for Japanese Biomedical and Behavioral Researchers at NIH (72402). PJD is supported by a NIH Oncology Research Training Grant (5T32CA009357).

## Authorship

Contribution: RS and YMS conceived and designed the study; RS, HNB, RR, PJD, PS, SBA and AVM developed the methodologies; RS, NWZ, ZHL, HNB, SS, WH, PS, NKK, SBA, AH and AVM acquired the data; RS, NWZ, ZHL, HNB, RR, PJD, PS, RK, AVM, CIKC, ST, EC analyzed and interpreted the data; RS, NWZ, ZHL, HJ, MPM, CAP, MDG, ADP, WZ, FJG, ES, CAL, and YMS supervised the study and wrote the manuscript; and all authors edited and provided input to the manuscript.

**Conflict-of-interest disclosure:** The authors declare no competing financial interests.

