## Supplementary figures and images for "Fatty acids in the tumor microenvironment reprogram neutrophils to induce immunosuppression via adenosine"

### Figure S1. Myeloid cells in the tumor microenvironment display immunosuppressive gene signatures (related to Figure 1).

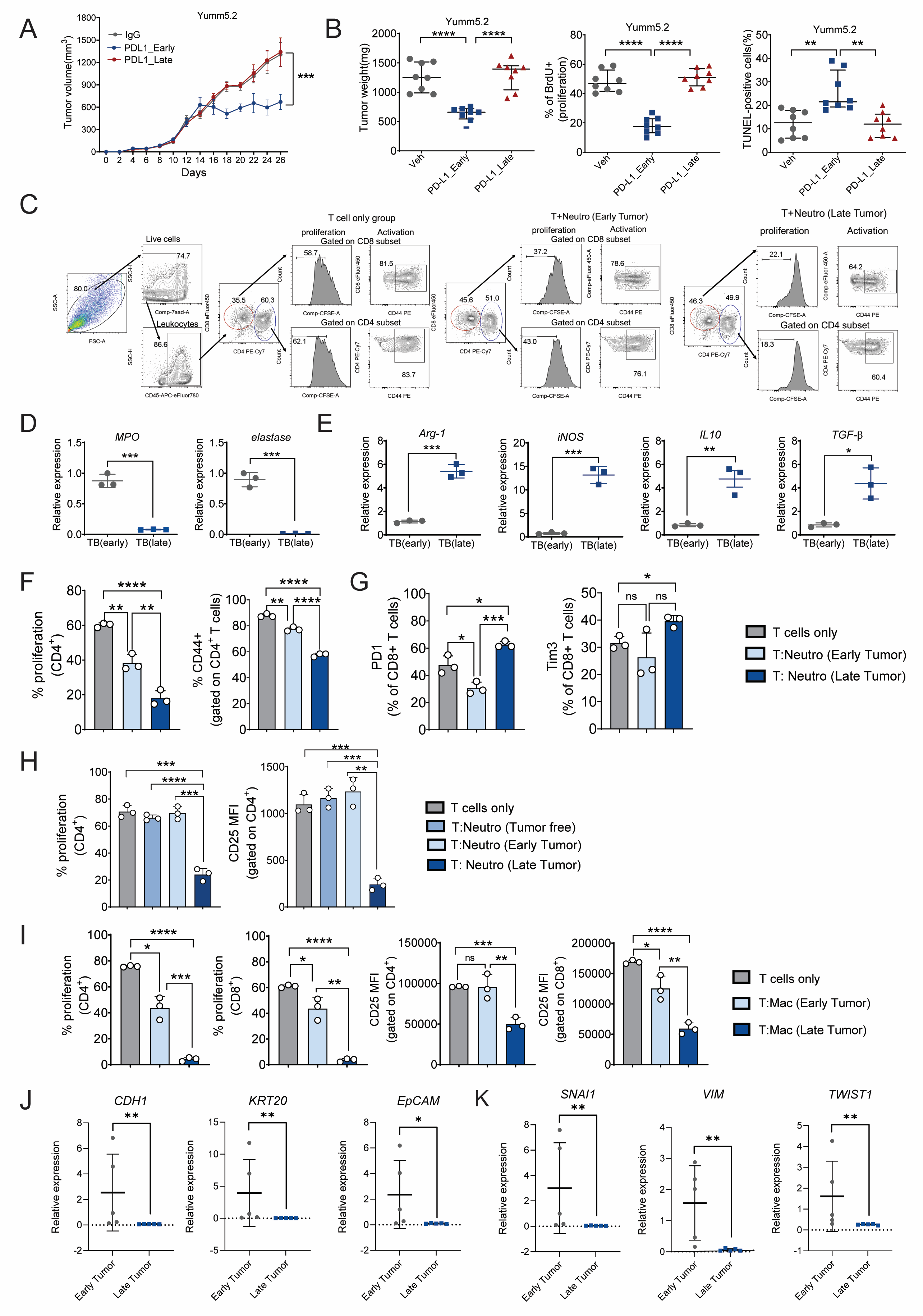

### Figure S2. TIF and serum from advanced tumor-bearing mice induce immunosuppression through neutrophils (related to Figure 2).

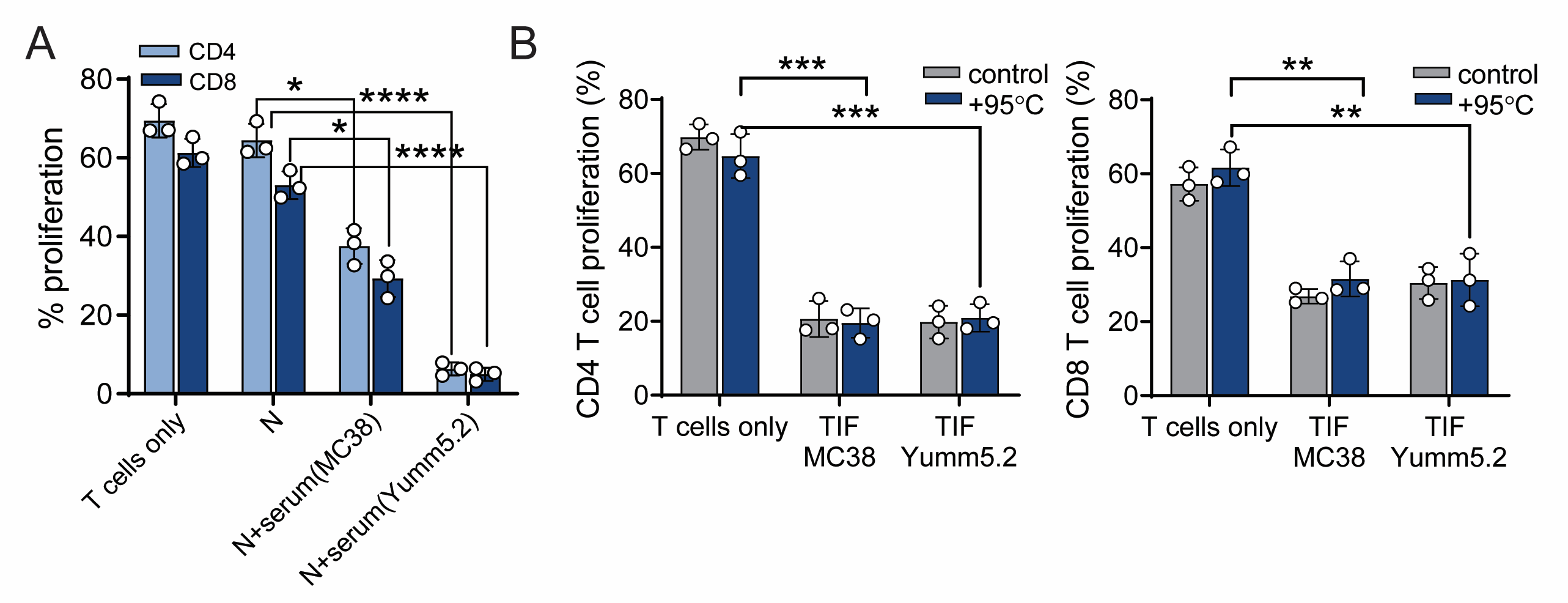

### Figure S3. Single-cell sequencing revealed heterogeneity in late-stage tumor neutrophils (related to Figure 3).

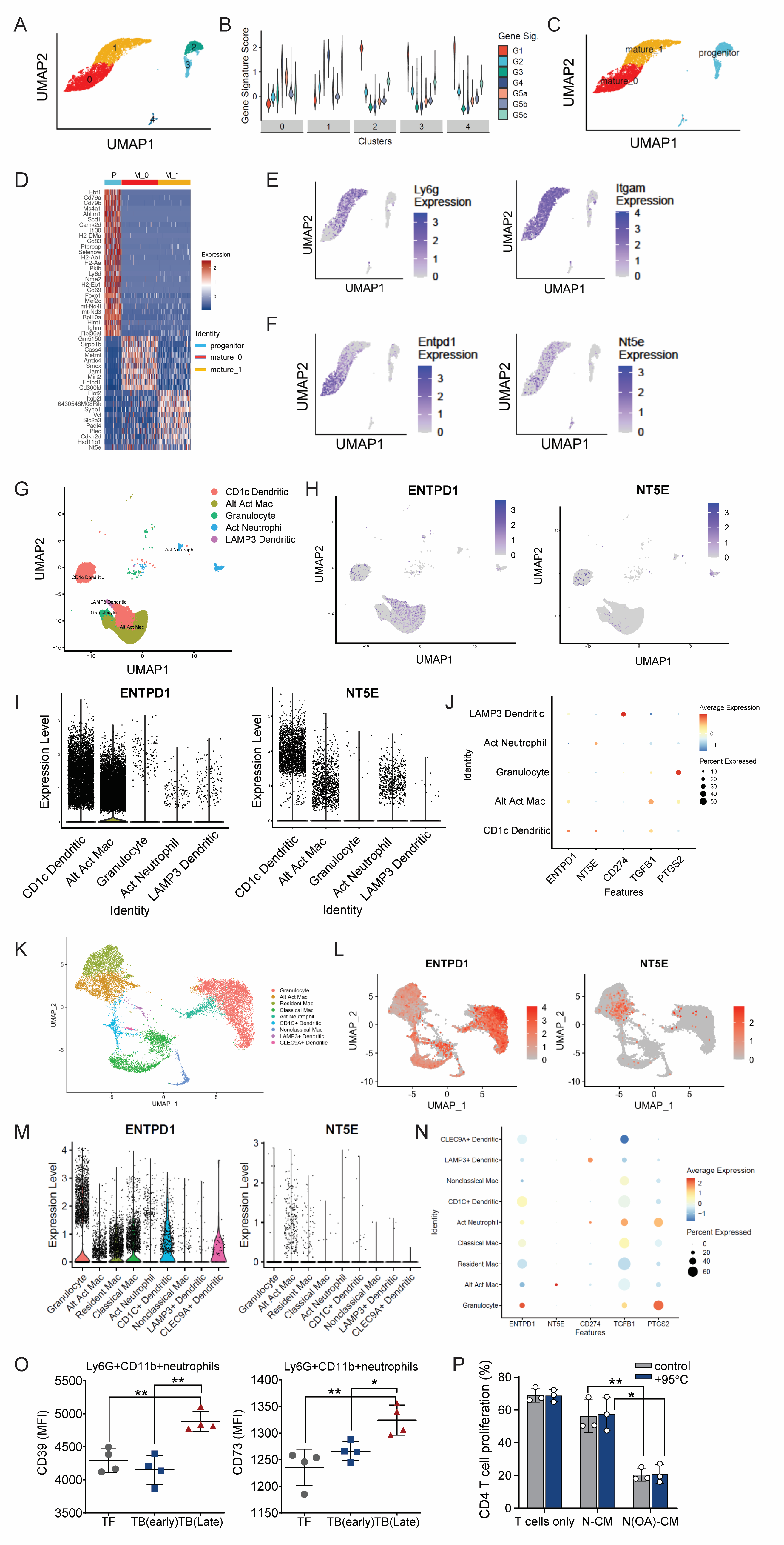

### Figure S5. AB680 and AB928 abolishes the suppressive effects of fatty acid-induced neutrophils on T cells (related to Figure 5).

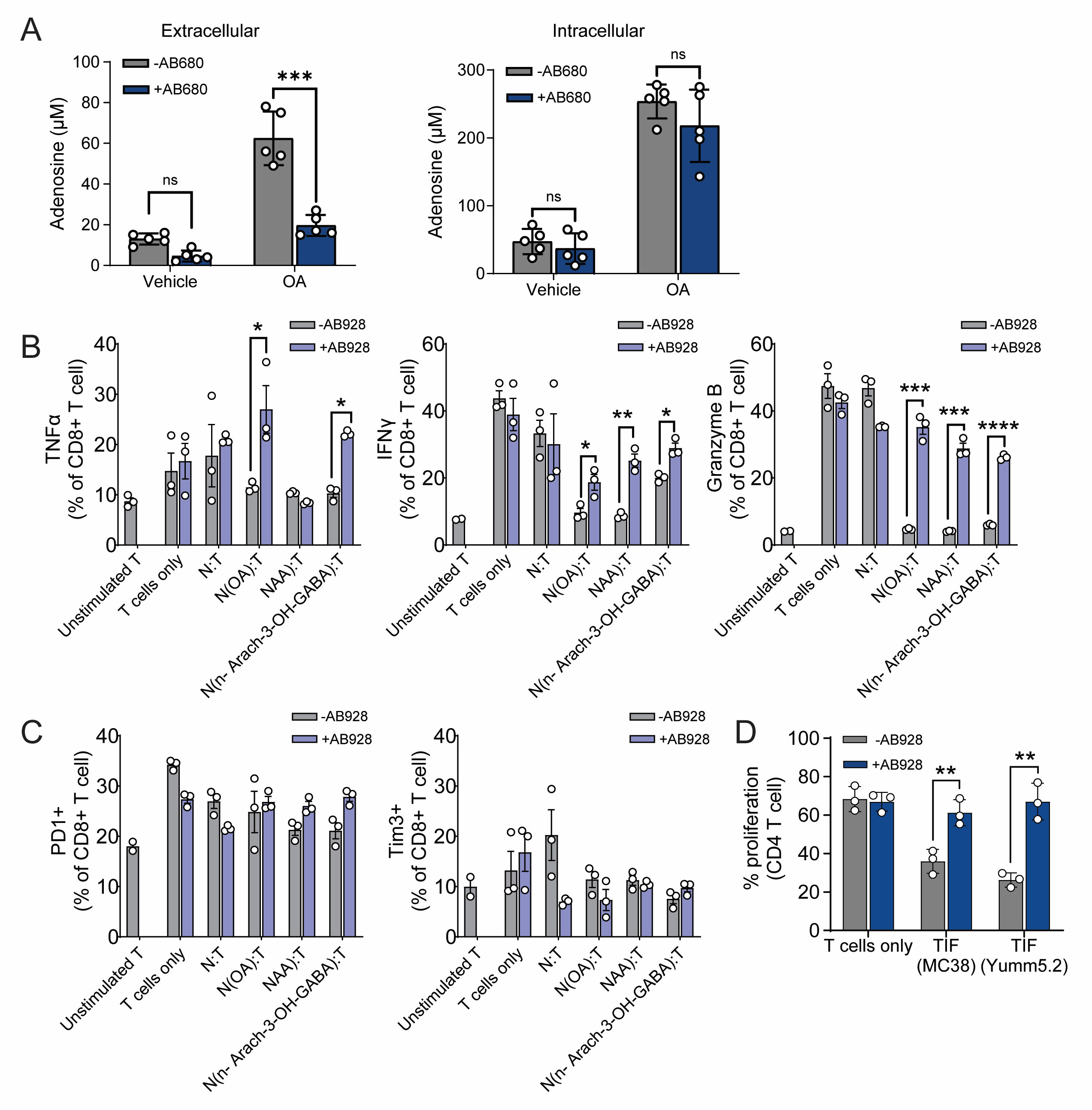

### Figure S6. AB928 restored the T-cell killing capacity in vivo (related to Figure 6).

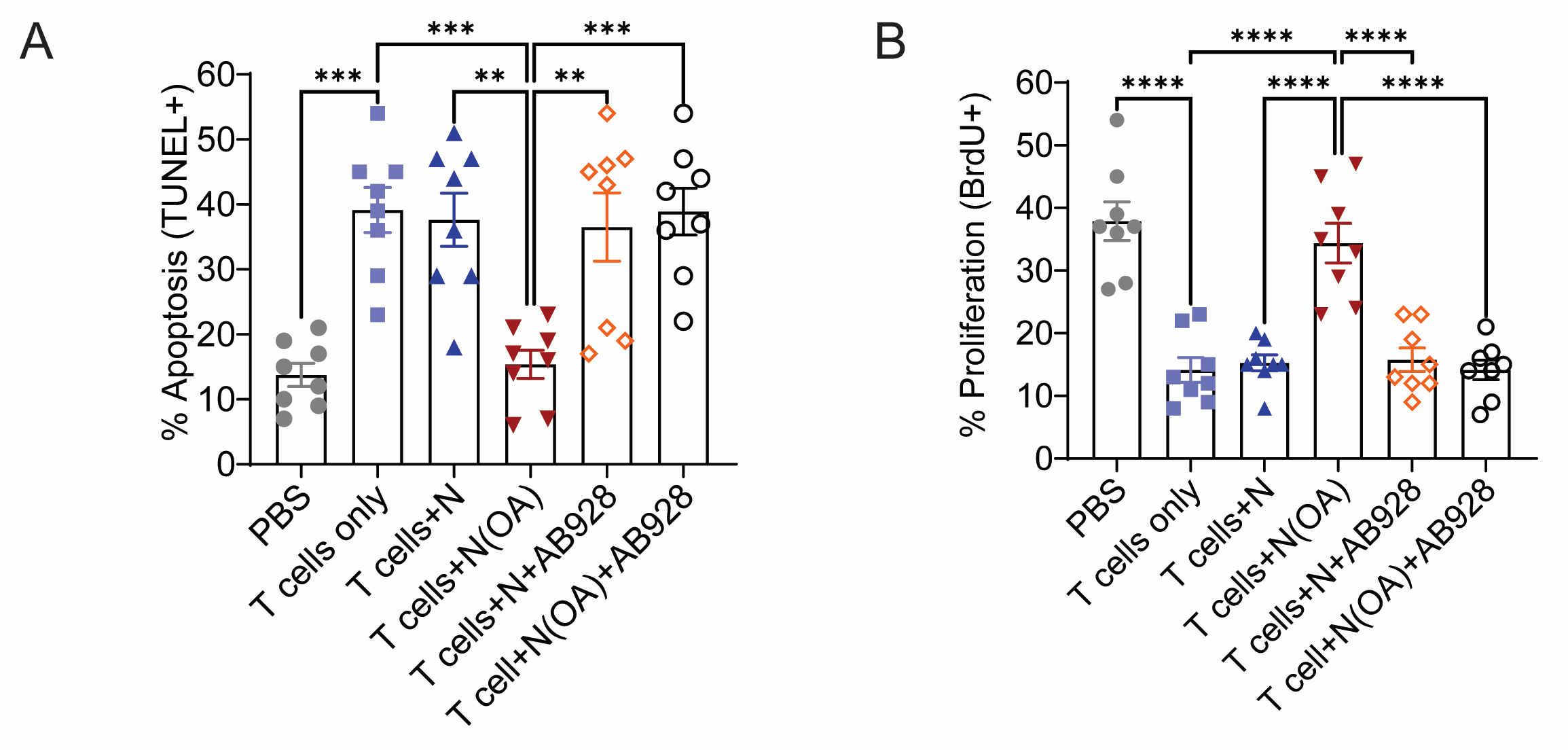

### Figure S7. Myeloid cells display high expression of adenosine signatures in an advanced autochthonous CRC mouse model.

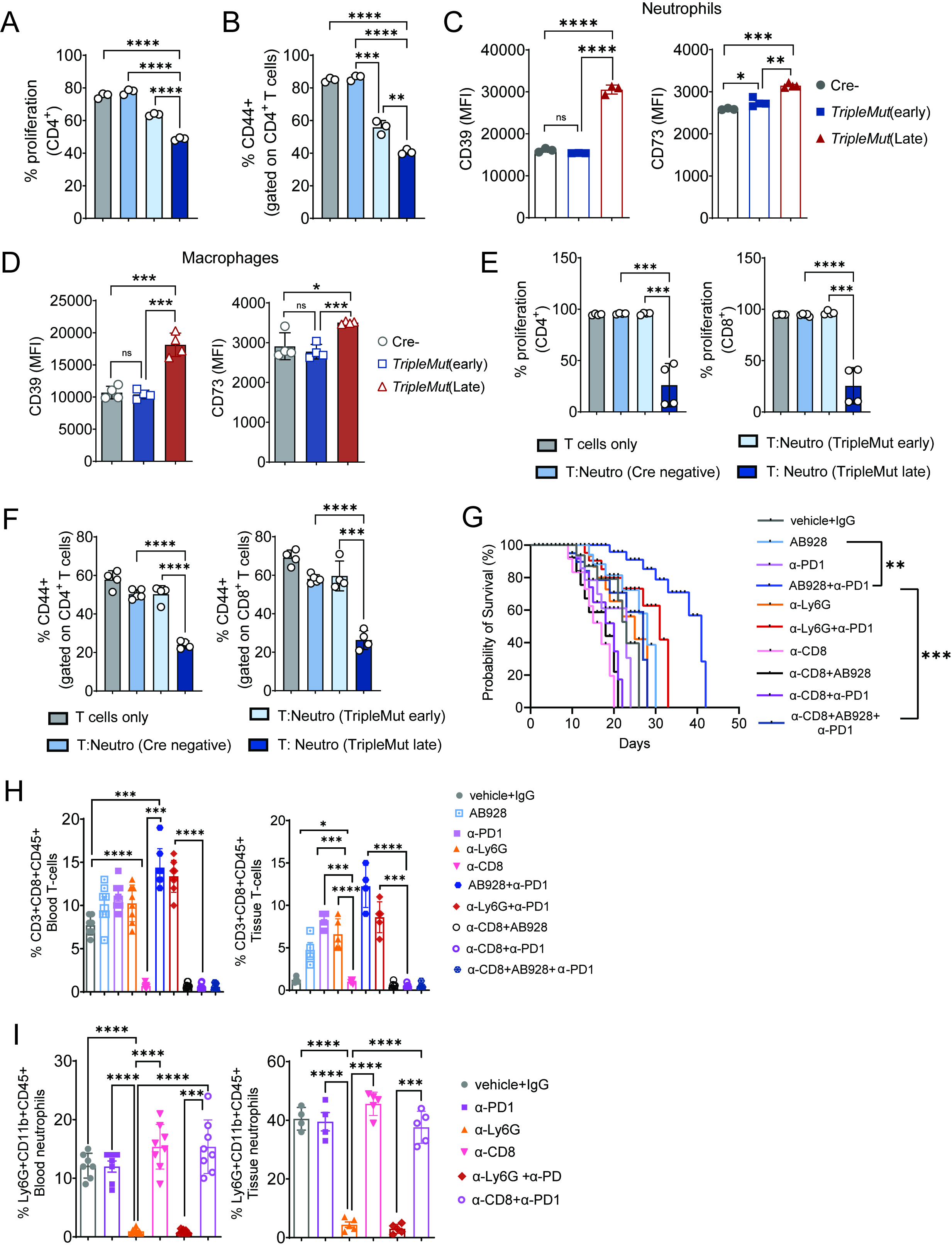

### Supplemental Data 1

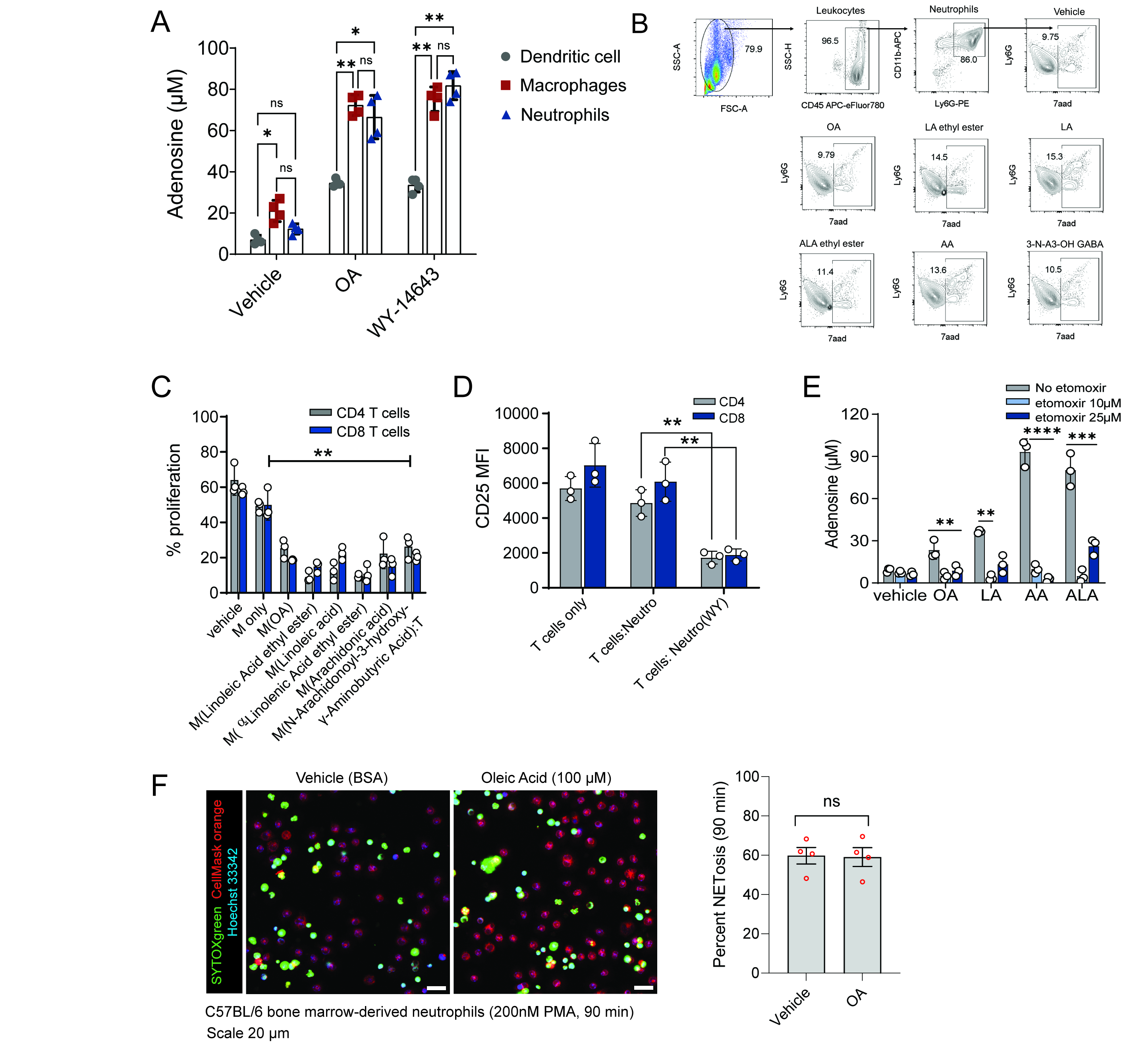
